# Brain single-cell transcriptional responses to bexarotene-activated RXR in Alzheimer’s disease model

**DOI:** 10.1101/2025.07.25.666575

**Authors:** Carolina Saibro-Girardi, Yi Lu, Nicholas F. Fitz, Daniel P. Gelain, Iliya Lefterov, Radosveta Koldamova

## Abstract

Pharmacological activation of brain Retinoid X Receptors (RXRs) enhances cognition and facilitates amyloid-beta (Aβ) clearance in Alzheimer’s disease (AD) mouse models, partly by upregulating Apolipoprotein E (*Apoe*), a major AD genetic risk factor. However, the specific cellular contributions to these effects are unclear. Here, we used single-cell transcriptomic profiling to investigate cell subpopulation-specific responses to bexarotene, an RXR agonist, in APP/PS1 mice. Our analysis revealed that bexarotene activated cholesterol biosynthesis and lipid metabolism transcriptional programs in homeostatic astrocytes and oligodendrocytes. Astrocytes also upregulated neurodevelopmental genes, while oligodendrocytes and endothelial cells showed enhanced protein folding and cellular growth pathways. Bexarotene further modulated immune responses, promoting Aβ-responsive signatures in disease-associated microglia and reactive astrocytes, while dampening pro-inflammatory responses in homeostatic microglia and endothelial cells. Furthermore, *Apoe* expression was significantly elevated across multiple cell types, especially in microglia and oligodendrocytes. Cell-cell communication analysis highlighted increased astrocyte-centered signaling, with APOE-driven pathways emerging as a prominent mediator. These findings clarify the cell-specific complexity of RXR-mediated regulation and underscore APOE as a central mediator of bexarotene’s neuroprotective effects. This study provides mechanistic insights into RXR-targeted interventions, and supports APOE-associated pathways as promising therapeutic targets in AD.

## Background

Alzheimer’s disease (AD) is the leading cause of dementia, accounting for up to 80% of all cases (1). It is characterized by extracellular β-amyloid (Aβ) plaques, Tau-containing intracellular neurofibrillary tangles, and neuroinflammation. Late-onset AD is believed to result from complex interactions between environmental and genetic risk factors, with *APOE* variants encoding apolipoprotein E (APOE) being the primary genetic contributor. APOE impacts amyloid deposition by interacting with Aβ, regulating its aggregation, clearance, and cell degradation in an isoform-dependent manner (2). Additionally, APOE-containing lipoproteins transport cholesterol from glial to neuronal cells (3). Beyond APOE dysfunction, AD involves metabolic disruptions (4), increased unfolded protein response (UPR) linked to Tau and Aβ (5), vascular dysfunctions (6), brain demyelination triggered by neuronal cell death (7), and impaired adult neurogenesis (8).

Targeting brain nuclear receptors (NR) has shown promise in AD mouse models (9, 10). NR are ligand-activated transcription factors regulating gene expression involved in energy metabolism, development, cell growth, and survival in response to lipophilic molecules. Retinoid X Receptors (RXR) are NR that act as obligatory heterodimerization partners for various family members, including Retinoic Acid Receptors (RAR), Thyroid hormone receptor, Liver X Receptors (LXR), and Peroxisome Proliferator-Activated Receptors (PPAR). RXR activation acts primarily through permissive dimers, where RXR agonists alone can drive transcriptional activity. In the brain, the permissive RXR partners mainly include LXR, linked to oxysterol responses, and PPAR, related to fatty acid (FA) responses, along with orphan NR. While endogenous RXR ligands such as 9-cis-retinoic acid (9-cis-RA) and 9-cis-13,14-dihydroretinoic acid have been identified, the physiological roles of RXR activation remain debated. Nevertheless, synthetic RXR-selective agonists have been developed, leading to numerous studies on their regulatory influence over complex gene networks (11).

Bexarotene, a synthetic RXR-selective agonist and an FDA-approved drug for cutaneous T cell lymphoma, has been extensively studied for its AD potential. It has shown significant cognitive and memory improvement in mice carrying familial AD mutations (12–14) and AD-associated APOE isoforms (15, 16). Bexarotene also facilitates soluble Aβ clearance, though its effect on plaque load remains unclear (16, 17). Studies, including our own, demonstrate that activated RXR-controlled networks influence brain immune response (18, 19), metabolism (20, 21), neurodevelopment (22–24), and myelination (25). At least partially, RXR-linked neuroprotective effects in AD are connected to APOE biology (9). However, these processes involve multicellular responses in the brain, and the mechanisms by which RXR activation affects different brain cell types remain unclear.

Therefore, our goal was to investigate the transcriptional events triggered by bexarotene in discrete brain cell populations. Using APP/PS1dE9 double-transgenic mice expressing APP and PS1 mutations (referred to as APP/PS1), we employed single-cell transcriptomics to achieve cell-level resolution *in vivo*. Our findings demonstrate that bexarotene-driven RXR activation promotes *Apoe* expression and regulates lipid metabolism, neurodevelopment, protein folding, and immune responses in a cell subtype-specific manner.

## Methods

### Animals

APP/PS1 mice were purchased from The Jackson Laboratory (USA), featuring the mutations APP K670_M671delinsNL (Swedish) and PSEN1:deltaE9. Cortical amyloid plaques begin to develop at three months of age and increase in size and number as the mice age. For experiments, animals were bred in-house as heterozygotes. Experimental mice were housed in groups of five females or four males per cage. Animals were littermates and maintained on a 12-hour light/dark cycle with *ad libitum* food and water. Both male and female mice were used in the experiments. All animal procedures adhered to the Guide for the Care and Use of Laboratory Animals from the United States Department of Health and Human Services, and were approved by the University of Pittsburgh Institutional Animal Care and Use Committee. All experiments were conducted in compliance with the ARRIVE guidelines.

### Mice treatment

We randomly assigned eight five-month-old APP/PS1 mice to either the Bexarotene treatment group (100 mg/kg/day bexarotene) or the Control group (vehicle treatment), ensuring a balanced distribution by age and sex. Bexarotene (Alfa Aesar) was prepared in a vehicle solution of corn oil containing 1% DMSO on the day of treatment. The animals received 10 ul/g of either bexarotene or vehicle solution by oral gavage for ten consecutive days (Table S1) (16, 18).

For euthanasia, mice were anesthetized with Avertin (1.25% tribromoethanol, 2.5% 2-methyl-2-butanol, 250mg/kg IP administration). Blood was collected via cardiac puncture of the right ventricle with EDTA-treated syringes followed by transcardial perfusion with 20mL of PBS. One brain hemisphere was dissected and immediately processed for brain dissociation and subsequent single-cell analysis.

### Neural tissue dissociation

We immediately processed freshly dissected mouse brain hemispheres, after removing the cerebellum, olfactory bulb, and subcortex, using the Adult Brain Dissociation Kit (Miltenyi Biotec) for mouse and rat, with slight modifications to the manufacturer’s protocol. Briefly, neural tissues were dissected into small pieces in HBSS and tissue pellets were incubated with enzyme mix P/Z (containing 0.125% 2-mercaptoethanol) for 15 min at 37°C, with gentle rotation every 2 min. Next, enzyme mix Y/A was added to the samples and the tissues were then mechanically dissociated by gentle pipetting and incubated for 10 min at 37°C with gentle rotation every 2 min. Samples were mechanically dissociated and incubated again for another 10 min at 37°C. The resulting suspensions were filtered through moistened 40μm cell strainers (Corning) and washed with HBSS (1:5). We then removed tissue debris using Debris Removal Solution (Miltenyi Biotec), following manufacturer’s instructions. Cell pellets were washed once using 0.04% BSA-DPBS buffer and filtered through 40μm Flowmi cell strainers (Bel-Art) before cell counting under the microscope using Trypan Blue. Cell viability ranged from 85 to 95% (Table S1).

### Single-cell transcriptomics

Droplet-based scRNA-seq library generation was prepared using Chromium Next GEM Single Cell 3’ Kit v3.1 and Chip G Single Cell Kit (10x Genomics) according to the manufacturer’s instructions. Immediately after tissue dissociation, Control-and Bexarotene-derived single-cell suspensions (four samples each) were used for GEM generation and barcoding in Chromium Controller (10x Genomics), with 10,000 cells as targeted recovery for each sample (Table S1). After cleanup, 25-75 ng of barcoded cDNA was amplified for library construction, and Dual Index Kit TT Set A (10x Genomics) was used for sample indexing. Library quality control was checked by Bioanalyzer High Sensitivity DNA kit (Agilent) (Table S1) and sequencing was performed on Illumina NovaSeq SP PE100 (MedGenome Inc.) targeting 400M reads per library.

### Data preprocessing

Single-cell barcoded reads were demultiplexed and aligned to the mouse reference genome (GRCm38) using Cell Ranger pipeline v7.0 (10x Genomics). The matrices of unique molecular identifiers (UMIs) generated by Cell Ranger were read into R v4.2.0 and analyzed using Seurat v4.2.1 (26), as previously described (27). Before processing, approximately 6.000 cells were sampled per library. The initial cell quality control step filtered only the cells with (i) unique feature counts > 200 and < 5000, (ii) total counts < 50000, and (iii) percentage of mitochondrial gene counts < 10, which resulted in a matrix with total 23,745 genes across 37,634 cells. SCTransform (28) was employed for feature normalization and scaling as well as variable feature analysis. We performed principal component (PC) analysis using variable features (RunPCA) followed by unsupervised clustering analysis (FindNeighbors using top 10 PCs, and FindClusters with 0.25 resolution). Uniform Manifold Approximation and Projection (UMAP) analysis was employed for dimensional reduction and data visualization (RunUMAP using top 10 PCs).

### Differential expression analysis

Clusters’ differential expression was performed using MAST v1.22.0 (29) within Seurat FindMarkers, then manually annotated based on the expression levels of cell type-specific gene sets (Table S1) (30–33). Next, selected cell types were further analyzed as subsets and similarly pre-processed. The mean expression of cell type-specific gene sets was added to the matrices using AddModuleScore, and an additional cell filtering step removed subclusters or cells expressing non-selective gene sets. Choroid-plexus epithelial cell gene *Ttr* and hemoglobin genes (*Hb*-) were disregarded in further analysis. Cell type-specific differential analysis between Bexarotene and Control groups was performed using MAST testing within Seurat. Differentially expressed genes (DEG) were defined according to adj. p-value < 0.05 and either pct.1 (fraction of Bexa-expressing cells) or pct.2 (fraction of Ctrl-expressing cells) > 0.2.

### Gene ontology and pathway analysis

Functional annotation of gene ontologies was performed using DAVID v6.8 web-tool (34), Cluster Profiler v4.6.2 gene set enrichment analysis with gseGO function (35), and Pathview for construction of pathway maps (36).

### Cell communication inference

We used CellChat v2 (37) to infer, analyze, and visualize cell-cell communication networks. The analysis input was a merged, preprocessed gene expression matrix from the subclusters Astro– Homeostatic, Astro–Reactive, Micro–Homeostatic, Micro–DAM, Oligodendrocytes and Endothelial–Capillary/Arteriole for each condition. We preprocessed and analyzed individual Control and Bexarotene matrices with CellChat to separately compute communication probabilities among cells and create independent cellular communication networks using computeCommunProb (with “trimean” as the method). We selected secreted signaling, ECM-Receptor, and Cell-cell contact pathways from CellChatDB, a manually curated ligand-receptor interaction database, for analysis.

Next, the networks were merged to compare Bexa vs. Ctrl. The differential number of interactions and the differential interaction strength were inferred using the functions compareInteractions and netVisual_diffInteractions. Interaction strength represents the computed communication probability, derived from two centrality measures: outdegree and indegree with weights. We assessed overall and cell pair-specific changes in signaling pathways by analyzing information flows, which are the sum of communication probabilities among all cell group pairs in the inferred network. We used the rankNet function, considering weights as measure. Finally, we evaluated changes in incoming and outgoing signaling pathways for specific cell populations using net analysis functions (netAnalysis_signalingRole_scatter and netAnalysis_signalingChanges_scatter).

### Statistical analysis

Sample sizes are indicated in the legends and correspond to biological replicates. Power analysis was performed to estimate the number of animals (two groups, t test, G*Power v3.1) with individual estimated experimental effect size, alpha =0.05, and 95% power. Statistical analyses referring to differential expression results were performed in R v4.2.0 using MAST v1.22.0 testing within Seurat and significance was determined as Bonferroni-corrected p-value <0.05 (adj.p-val). Cluster distribution among libraries was analyzed on GraphPad Prism v8.0.2 using two-way ANOVA and significance for the effect of treatment was determined as p-value <0.05. DAVID v6.8 GO analysis used Fisher’s exact test, and cell communication inference (CellChat v2) used paired Wilcoxon test, both considering p-value <0.05. Unless otherwise indicated in figure legends, results are reported as means with error bars as standard error (SEM).

## Results

### Single-cell transcriptomics of APP/PS1 mouse brains identify glial, neuronal, and vascular cell populations

To get further insight into the molecular mechanisms underlying the therapeutic effects of brain RXR activation, we treated five-month-old APP/PS1 mice with bexarotene (Bexa) or vehicle (Ctrl) through oral gavage for ten consecutive days, following previous studies (16, 18). At this stage, APP/PS1 mice exhibit early AD pathophysiology (38). The dissected cortical plate was enzymatically dissociated to generate cell suspensions for single-cell RNA sequencing (scRNA-seq) workflow (Figure 1A). After cell quality control and filtering (Figure S1A-B), transcriptional data from 37,634 high-quality cells was preprocessed (28). Dimensional reduction followed by clustering analysis (26) identified 19 discrete cell populations across all samples (Figure 1B). Cluster differential expression analysis (Figure S1C) and evaluation cell type-specific gene sets (Figure S1D) enabled precise cell population identification (Figure 1C).

**Figure 1.**
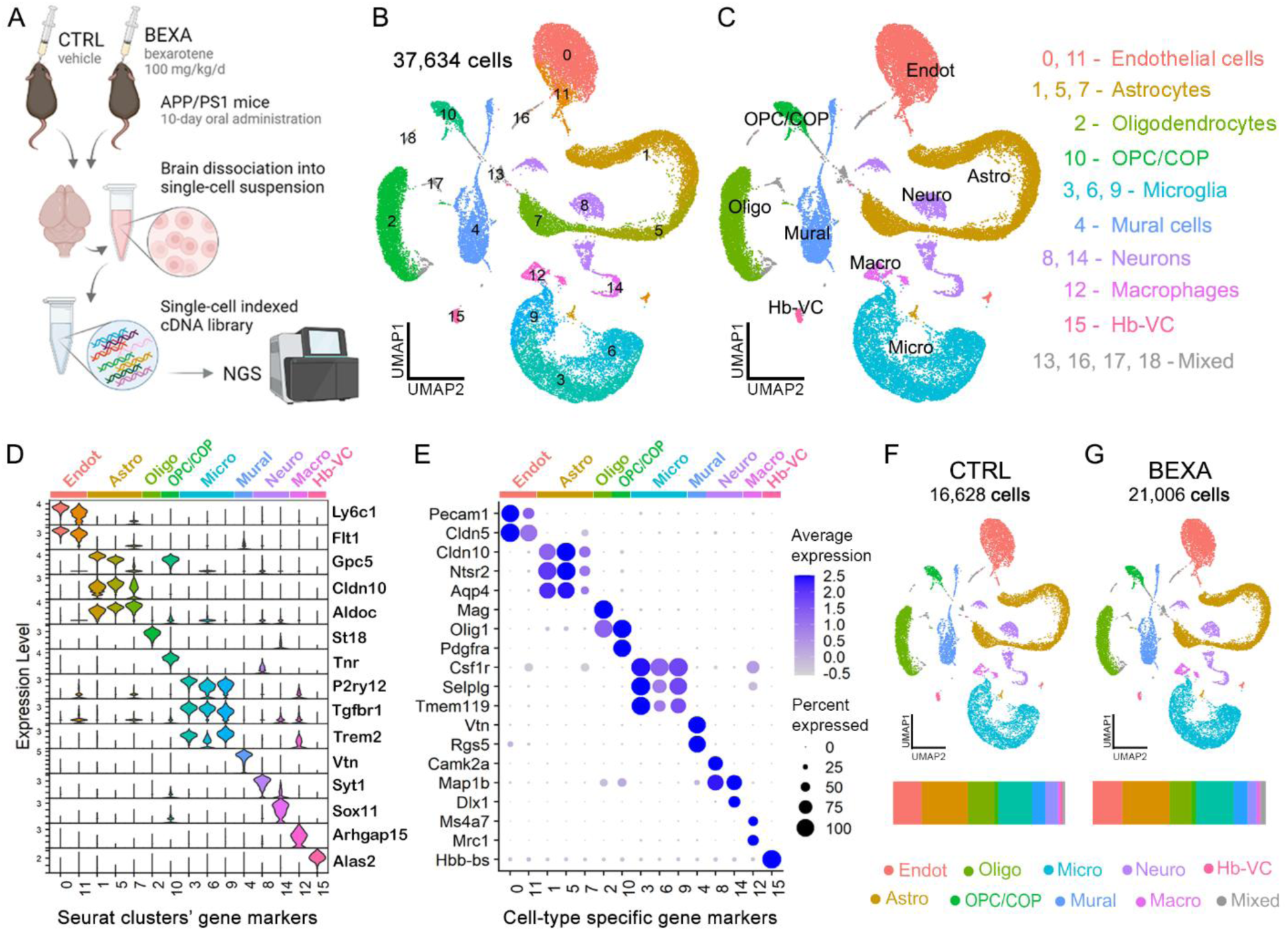
scRNA-seq data of bexarotene-treated APP/PS1 mice brains. (A) Study design and scRNA-seq workflow. APP/PS1 mice (5 mo., male/female) were treated with bexarotene (Bexa, 100 mg/kg/d) or vehicle (Ctrl, corn oil with DMSO 1%) for 10 days through oral administration. Next, the mice brains were dissected and one hemisphere (cerebellum, olfactory bulb and subcortex removed) was dissociated into single cell suspensions. Brain-derived cells were immediately used for single-cell RNA-seq workflow using 10X Chromium platform and next generation sequencing (NGS). (B) UMAP dimensional reduction of 37,634 high quality cells detected. Cell ranger software was used for alignment and demultiplexing, and Seurat-based SCTransform normalization, PCA analysis and unsupervised clusterization found 19 clusters. (C) UMAP plots grouped by cell type, identified after clusters’ DE analysis using MAST. We identified astrocytes (Astro, 10,155 cells), microglia (Micro, 7,820), oligodendrocyte lineage (Oligo and OPC/COP, 6,159), neuronal lineage (Neuro, 2,296), blood vessel cells (Endot and Peric, 8,421) and brain monocyte/macrophages (Macro, 636). Clusters 13, 16, 17 and 18 comprised mixed cell populations or doublets. (D) Expression level of clusters’ gene markers and (E) average expression plus percent of cells expressing examples of cell type markers. (F) UMAP plots grouped by cell type split by group treatment, with control (Ctrl, 16,627 cells) and bexarotene (Bexa, 21,006 cells). The bars at the bottom show the distribution of cell type populations sampled for each respective group.

Figure 1D illustrates the expression levels of top marker genes for each cluster, and Figure 1E highlights key cell type-specific genes used for annotation. The glial population included astrocytes expressing *Cldn10, Ntsr2*, and *Aqp4* (clusters 1, 5, and 7) and microglia expressing *Csf1r, Selplg*, and *Tmem119* (clusters 3, 6, and 9). The oligodendrocyte lineage featuring *Olig1* expression included mature oligodendrocytes (cluster 2) expressing *St18* and *Mag* markers of myelinating cells, and immature cells (cluster 10) expressing *Tnr* and *Pdgfra*. The dataset also included brain blood vessel cells: endothelial cells (EC) expressing *Flt1* and *Cldn5* (clusters 0 and 11) and mural cells expressing *Vtn* and *Rgs5* (clusters 4 and 16). Neurons accounted for approximately 6% of the dataset, with clusters 8 and 14 encompassing a heterogeneous neuronal population. Minor brain cell populations included perivascular/parenchymal macrophages expressing *Ms4a7* and *Mrc1* (cluster 12), and choroid plexus epithelial and ependymal cells (cluster 13) expressing *Prlr* and *Tmem212*, respectively (Figures 1D-E and S1D). Clusters 13, 16, 17, and 18 comprised mixed cell populations or doublets (Figure S1D). Comparison between Ctrl and Bexa datasets showed similar cell quality control metrics (Figure S1B), with merging UMAP plots and comparable cell type-distributions, despite a higher number of sampled cells in the Bexa group (Figure 1F).

### Bexarotene activates lipid biosynthesis and brain development in homeostatic astrocytes

Astrocytes represented the largest cell population in the APP/PS1 mouse brain libraries. To focus on astrocyte-specific transcriptional changes, we isolated the astrocyte clusters (1, 5, and 7), reprocessed the data, and filtered out cells expressing unspecific genes. This yielded 10,155 high-quality astrocytes across Ctrl and Bexa groups. Re-clustering identified three discrete subpopulations (Figure 2A). Astro0 exhibited high expression of genes linked to astrocyte cell adhesion (*Lsamp*), as well as genes linked to brain development, synaptic transmission, and axon guidance, resembling homeostatic astrocytes (39) (Figures 2B and S2A). Astro1 was enriched for *Aldoc* (Aldolase C) and other genes associated with metabolic processes and redox homeostasis, indicative of astrocyte reactivity (40). Astro2 selectively expressed *Crym* (Crystallin) alongside high expression of other proliferation-associated genes, suggesting a population encompassing proliferative astrocytes (Figures 2B and S2A). Astrocyte subcluster distribution was similar between Ctrl and Bexa (Figure S2B).

**Figure 2.**
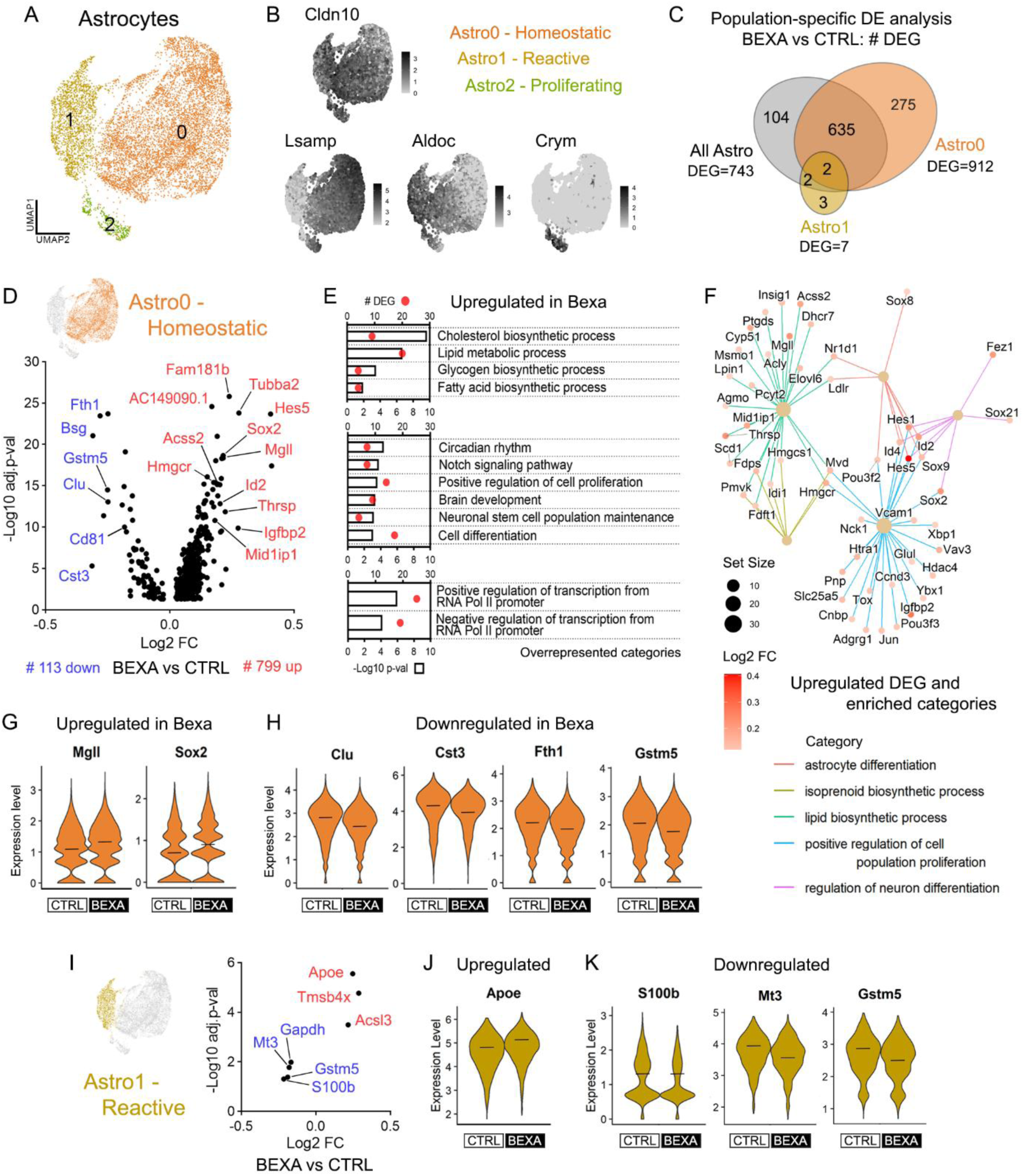
Bexarotene-mediated transcriptional responses in homeostatic and reactive astrocytes. (A) UMAP plot of the 10,002 astrocytes revealing 3 subclusters: Astro0 – Homeostatic, Astro1 – Reactive, and Astro3 – Proliferative. (B) Feature plots of subclusters’ gene markers: Cldn10 (all Astro), Lsamp (Astro0), Aldoc (Astro1), and Crym (Astro3). Scales indicate the expression level. (C) Venn diagram of Bexa vs Ctrl DE analysis with total, exclusive, and shared DEG numbers in astrocyte populations. Tests considered Astro0, Astro1, or all astrocyte cells. DEGs display adj.p-val<0.05 and are expressed in >20% of Ctrl or Bexa cells. (D) Volcano plot of 799 upregulated and 113 downregulated DEG in Bexa vs Ctrl comparison within Astro0 – Homeostatic astrocytes. (E) Overrepresented GO terms (biological process) associated with the list of log2FC>0.1 upregulated genes in Bexa group within Astro0. Bars indicate -log10 p-val and dots indicate DEG count. (F) Category netplot of upregulated DEG in Bexa group within Astro0, connected with the categories found in GSEA analysis. Scales indicate the gene set size and DEG log2FC. (G-H) Expression level of upregulated (Mgll, Sox2) and downregulated (Clu, Cst3, Fth1, Gstm5) genes, respectively, within Astro0 cells. (I) Volcano plot of DEG in Bexa vs Ctrl comparison within Astro1 – Reactive astrocytes. (J-K) Expression level of upregulated (Apoe) and downregulated (S100b, Mt3, Gstm5) genes, respectively, within Astro1 cells. Crossbars indicate mean values, and the expression levels are displayed as SCTransform corrected UMI counts. Sample size: Bexa = 4,305, Ctrl = 3,280 (Astro0); Bexa = 1,154, Ctrl = 879 (Astro1).

Next, we performed Bexa vs. Ctrl differential expression (DE) analysis considering either the total astrocyte population or selective subclusters. Figure 2C shows that most differentially expressed genes (DEG) were concentrated in Astro0-Homeostatic astrocytes. DE analysis within Astro0-Homeostatic cells identified 912 DEG, 799 up- and 113 downregulated (Figure 2D and Table S2). Numerous genes involved in lipid metabolism were upregulated. The enriched GO term Cholesterol biosynthesis process included coding genes for enzymes in the mevalonate pathway (*Hmgcr, Pmvk, Mvd*, *Fdps, Fdft1, Cyp51, Msmo1, and Dhcr7*) (Figures 2D-E and S2C), which is controlled by both LXR/RXR and PPAR/RXR dimers (41). Likewise, the terpenoid backbone biosynthesis and steroid biosynthesis pathways were enriched (Figure S2D-E). For instance, the upregulated *Hmgcr* (Figure S2D) encodes the rate-limiting enzyme HMG-CoA reductase for cholesterol synthesis. The upregulation of lipid metabolic processes also included lipogenic genes (*Acss2, Mid1lip1*, *Thrsp*) and genes linked to FA biosynthetic process (*Mgll, Elovl6, Scd1*, *Ptgds*) (Figures 2D-F and S2C). The lipase-coding *Mgll* (Figure 2G) converts monoacylglycerols into free FA and glycerol, while LXR/RXR target genes *Elovl6* (ELOVL Fatty Acid Elongase 6) and *Scd1* (Stearoyl-CoA Desaturase 1) are crucial for FA elongation (42). These findings indicate that bexarotene stimulates cholesterol and FA biosynthesis in homeostatic astrocytes, vital for brain homeostasis, learning, and cognition (3).

Genes involved in brain development were also activated within Astro0 (Figure 2D-F). Examples include transcription factors (TF)-coding genes such as *Sox2* (Figure 2G)*, Sox9,* and *Sox21* from the SRY-box family, as well as *Hes5* (Figure S2C) and *Hes1* from the Hes family. The genes *Id2* (Figure S2C) and *Id4,* from the inhibitors of DNA-binding family, were also upregulated. These family of genes play key roles in the enriched terms Cell differentiation, Neuronal stem cell population maintenance, Positive regulation of cell proliferation, Notch signaling pathway and Circadian rhythm (Figure 2E). These TF participate in astrocyte differentiation and the regulation of neuronal differentiation (Figure 2F) and play fundamental roles in the stemness and differentiation of adult neural progenitor cells. The upregulated *AC149090.1,* encoding a phosphatidylserine decarboxylase, is recognized as an ageing clock in neural stem cells (NSC), involved in both neurodevelopment and metabolism (43). Given the transcriptional similarities between astrocytes and NSC (44), Astro0 possibly contains a mixture of mature astrocytes and immature glial cells, including adult stem cells. Furthermore, the numerous DEG linked with positive regulation of cell proliferation (*Igfbp2, Ccnd3, Sox2, Hes5, Id4, Jun*) indicate stimuli promoting the cell growth of non-reactive astroglia (Figure 2F). These findings align with our previous studies linking RXR activation with brain regeneration and the differentiation of both neurons and glial cells (22–24). Additionally, several TF were upregulated, as illustrated by the terms Positive and Negative regulation of transcription from RNA Pol II promoter (Figure 2E), indicating extensive secondary transcriptional events in homeostatic astrocytes.

The top downregulated genes by bexarotene included *Clu* (Clusterin), *Cst3* (Cystatin C), *Fth1* (Ferritin heavy chain 1), and *Cd81* (Tetraspanin) (Figure 2D and 2H), all of which are associated with the AD-associated astrocyte phenotype (39). The downregulation of the glutathione S-transferase coding gene *Gstm5* (Figure 2H) was observed in both Astro0 and Astro1. *Fth1* and *Gstm5* are important for oxidative stress responses, along with other downregulated genes (*Scara3, Cox4l1, Cox6c, Mgst1, Txnip, Prnp*) (Table S2). *Cst3, Clu,* and *Cd81* are involved in astrocyte immunoreactivity and cell proliferation, with Cystatin C and Clusterin being well-known players in Aβ trafficking and plaque-associated responses (Figure 2G) (32, 39). Conversely, *Abca1* (ATP-binding cassette transporter A1), an LXR target gene linked to lipid transport was also downregulated, though to a lesser extent (Table S2). Overall, our findings indicate that neuroinflammatory, stress-related, and AD-associated features are reduced in homeostatic astrocytes following bexarotene treatment.

### Reactive astrocytes display upregulated *Apoe* and suppressed stress-related genes under RXR activation

Reactive astroglia refers to the astrocyte population that undergoes cellular programs in response to a pathological context, leading to a range of reactive phenotypes (40). Among them, an AD-unique disease-associated astrocyte population has been reported, expressing high levels of genes related to amyloid metabolism and clearance (39). Within the reactive astrocytes recognized as Astro1 in the APPS/PS1 model, we identified fewer DEG induced by bexarotene compared to Astro0-Homeostatic (Figure 2C and Table S2). Astro1-Reactive showed 7 DEG in Bexa vs Ctrl (Figure 2I), 5 exclusive to Astro1 including the AD-linked *Apoe* (Figure 2J), *Acsl3* (Acyl-CoA Synthetase Long Chain Family Member 3) involved in lipid biosynthesis and beta-oxidation, and *Tmsb4x* (Thymosin beta-4) linked to cytoskeleton organization. *Apoe* expression is a fundamental feature of astroglia and, in AD-associated astrocytes, it is also linked to Aβ reactivity (45, 46). Its upregulation bexarotene-exposed Astro1 thus suggests increased Aβ reactivity in reactive astrocytes following the treatment.

Conversely, *S100b* (S100 Calcium Binding Protein B)*, Mt3* (Metallothionein 3), and *Gstm5* are also involved in astrocyte reactivity but were downregulated in Astro1 (Figure 2K). S100B, also downregulated in Astro0, is recognized as a protein expressed and released by astrocytes under brain injury, acting as a damage-associated molecular pattern (47). Together with the roles of *Mt3* and *Gstm5* activation in oxidative stress, our findings thus indicate reduced stress responses in reactive astrocytes of bexarotene-treated APP/PS1 mice.

### Bexarotene activates microglial *Apoe* and differentially modulates immune cell reactivity in homeostatic and DAM populations

Next, we analyzed microglial cell populations in the APP/PS1 mouse brain. Clusters 3, 6, and 9 were analyzed as a subset yielding final 7,567 high-quality cells. Re-clustering identified three distinct populations: Micro0-DAM1, Micro1-Homeostatic, and Micro2-DAM2 (Figure 3A). Disease-associated microglia (DAM) are a subset of brain immune cells found at neurodegeneration sites, playing protective roles through the activation of phagocytic, lysosomal, and lipid metabolism pathways (48). Cluster Micro1-Homeo displayed gene expression associated with homeostatic microglial functions, such as *Malat*, *Nav3* and *Tanc2* (Figure 3B and S3A). By contrast, both Micro0 and Micro2 displayed high levels of classical DAM genes like *Tpt1*, *Tyrobp*, *Trem2*, and *Apoe*, but the 511 cells of Micro2-DAM2 selectively expressed *Cst7* which is spatially correlated with amyloid plaques (27, 33, 49) (Figure 3B and S3A-B). Micro0-DAM1 also expressed relatively high levels of *Cst3, Aif1*, and *C1q* family of complement genes (Figure S3A). Comparing cluster distributions between treatment groups revealed equivalent proportions: 43% homeostatic (Micro1) and 57% DAM (Micro0 & Micro2) (Figure S3C). We merged Micro0 and Micro2 into a single DAM cluster and conducted Bexa vs Ctrl DE analysis across the entire microglial population, Micro-Homeo, and Micro-DAM. Compared to astrocytes, microglia exhibited fewer DEG: 66 genes in Homeo, 40 genes in DAM, and 47 genes when all microglia were considered (Figure 3C and Table S2). Notably, the gene sets affected in Micro-Homeo and Micro-DAM were largely distinct (Figure 3C-E). Figures 2F and 2G respectively compare the gene sets upregulated and downregulated by bexarotene in both microglial subpopulations.

**Figure 3.**
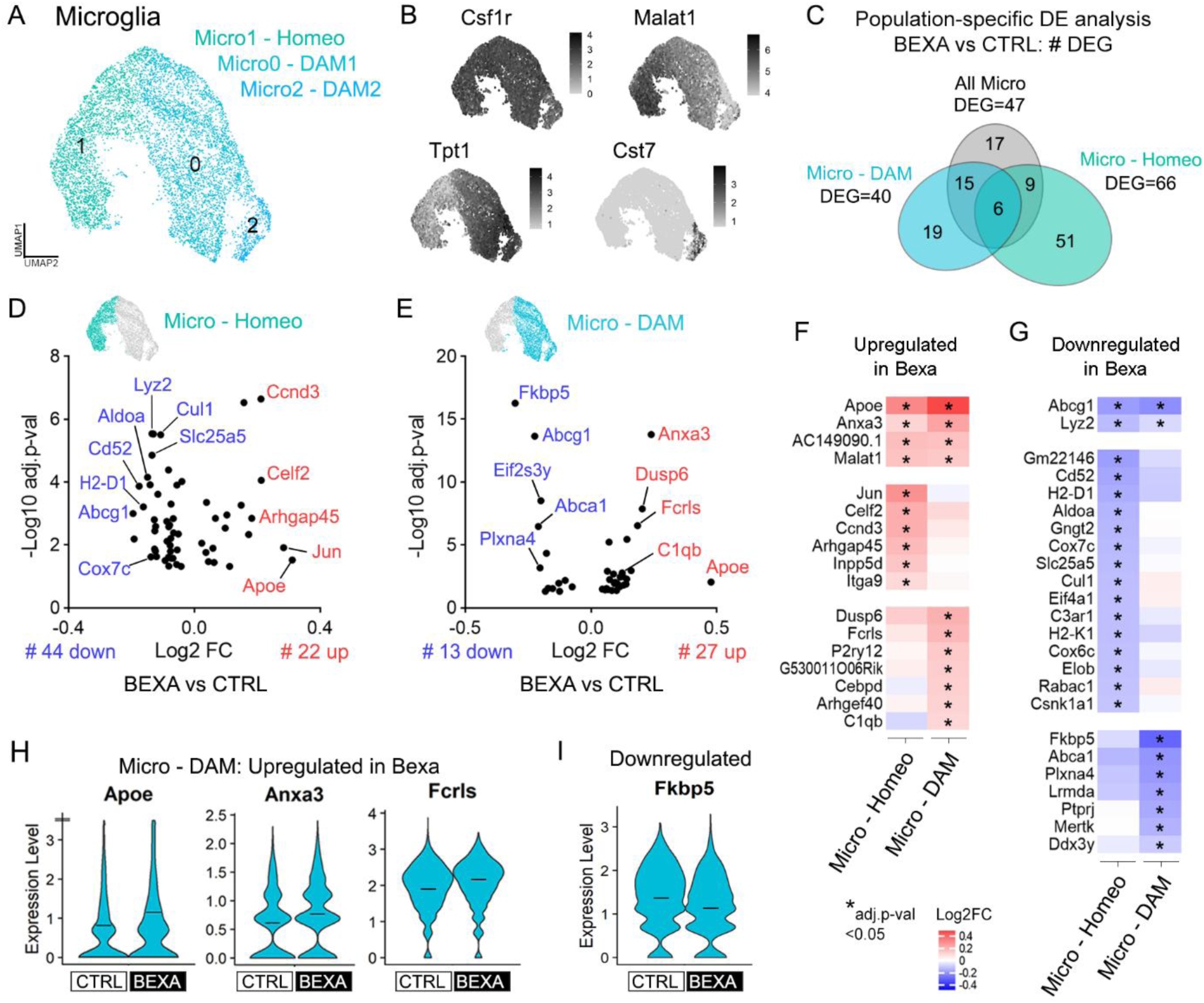
DAM and homeostatic microglia-specific transcriptional responses in APP/PS1 mice treated with bexarotene. (A) UMAP plot of the 7,106 cells showing Micro0, Micro1, and Micro2 subclusters. (B) Feature plots of subclusters’ gene markers: Csf1r (all Micro), Malat1 (Micro1, homeostatic), Tpt1 (Micro0, DAM), and Cst7 (Micro1, DAM). (C) Venn diagram of Bexa vs Ctrl DE analysis with total, exclusive, and shared DEG numbers in microglia populations. Tests considered the homeostatic (Micro1), DAM (Micro0 & Micro2), or all Micro populations. DEGs display adj.p-val<0.05 and are expressed in >20% of Ctrl or Bexa cells. (D-E) DE analysis within Micro – Homeo and Micro – DAM, respectively. Volcano plots of 22 up- and 44 downregulated DEG within Micro – Homeo, and of 27 up- and 13 downregulated DEG within Micro – DAM population. (F-G) Heatmaps showing Bexa vs Ctrl fold change of upregulated and downregulated DEG, respectively, in microglia subpopulations. DEG in both identities (top panels), in Micro – Homeo only (middle), and in Micro – DAM only (bottom). Scales indicate Bexa vs Ctrl log2FC, and * indicates adj.p-val<0.05. (H-I) Expression level of upregulated (Apoe, Anxa3 and Fcrls) and downregulated (Fkbp5) genes, respectively, within Micro – DAM population. Crossbars indicate mean values. Expression levels are displayed as SCTransform corrected UMI counts. Sample size: Bexa = 2,396, Ctrl = 1,662 (DAM); Bexa = 1,861, Ctrl = 1,187 (Homeo).

Within homeostatic microglia (Micro-Homeo), bexarotene upregulated 22 genes and downregulated 44 (Figure 3D). *Celf2* and *Jun* homeostatic genes were selectively upregulated within Micro-Homeo, as well as the cyclin-coding gene *Ccnd3* (Figure 3F). Conversely, bexarotene selectively downregulated the numerous DAM-associated genes (*Cd52, Aldoa, Cul1)*, histocompatibility genes (*H2-D2, H2-K1*), and cyclooxygenase-coding genes (*Cox7c, Cox6c*) (Figure 3G). These changes indicate that bexarotene enhances the surveilling and physiological features of homeostatic microglia.

Within the DAM population (Micro-DAM), bexarotene induced the activation of 27 genes and the downregulation of 13 (Figure 3E). *Apoe* and the microglial gene *Anxa3* (Annexin 3) were top bexarotene-induced genes within this population, also upregulated in Micro-Homeo but to a lesser extent (Figure 3F). *Apoe* (Figure 3H) is crucial for DAM phenotype acquisition, acting also via the APOE/TREM2 axis to direct lipoprotein-ligated Aβ for degradation by microglia (48). *Anxa3* (Annexin 3) (Figure 3H) regulates cell growth in immune cells and is found upregulated in activated microglia (50). In Micro-DAM, upregulated genes were also connected to the activation/control of MAPK signaling pathways (*Anxa3*, *Dusp6*, *Fcrls*) (Figure 3E-F). For instance, *Fcrls* (Figure 3H), an Fc receptor family member, interacts with opsonized immune complexes driving its phagocytosis. The complement gene C1qb was also upregulated by bexarotene within DAM cells (Figure 3E-F). These results indicate that bexarotene enhances immune responses and Aβ reactivity in DAM of APP/PS1 mice. A similar response has been reported in macrophages (18, 51), though we did not detect DEG in cluster 12 containing macrophages (not shown).

The top downregulated gene in Micro-DAM was *Fkbp5* (Figure 3G and 3I), encoding a peptidyl-prolyl isomerase with co-chaperone activity known to promote neuroinflammation and cytotoxicity through NF-κB activation (52). This finding supports the role of RXR activation in reducing neuroinflammation through LXR- and PPAR-mediated inhibition of NF-κB (41), and aligns with PPARα-induced suppression of FKBP5 in the APP/PS1 cortex (53). Lastly, *Lyz2* and the LXR target gene *Abcg1* were downregulated by bexarotene in both DAM and homeostatic microglia, while *Abca1* was specifically downregulated in Micro-DAM (Figure 3G). This indicates that Bexa-associated *Apoe* upregulation in microglia is functioning through ABCG1/ABCA1-independent immune pathways (54–57).

### Oligodendrocytes display upregulated cholesterol synthesis and protein folding gene expression following bexarotene treatment

We next analyzed the oligodendrocyte lineage subset, encompassing clusters 2 and 10. The final dataset comprised 5,898 high-quality cells, primarily mature oligodendrocytes alongside a population containing oligodendrocyte precursor cells (OPC) and differentiation-committed oligodendrocyte progenitors (COP) cells (Figure S4A) (30), with similar cluster distribution between treatment groups (Figure S4B). Oligodendrocytes expressed the marker of oligodendrocyte lineage *Olig1*, along with key myelination markers such as *Plp1*, *Mbp,* and *St18* (Figure 4A-B and S4A). Bexa vs. Ctrl DE analysis revealed 141 DEG in oligodendrocytes following bexarotene treatment (Figure 4C and Table S2). Of these, 79 genes were upregulated, with *Apoe* emerging as the top activated gene (Figure 4C-D). While APOE is extensively studied in astrocytes and microglia, it also plays a critical role in oligodendrocytes, as dysfunctional APOE biology is linked to oligodendrocyte cholesterol accumulation and impaired myelin formation (58). Another player in oligodendrocyte lipid dynamics, *Apod*, was also upregulated by bexarotene (Figure 4D). APOD lipoprotein is essential for cell lipid homeostasis, controlling lipids’ redox state across subcellular/extracellular locations (59). In oligodendrocytes, APOD further contributes to the compaction of the extracellular leaflet of myelin (60).

**Figure 4.**
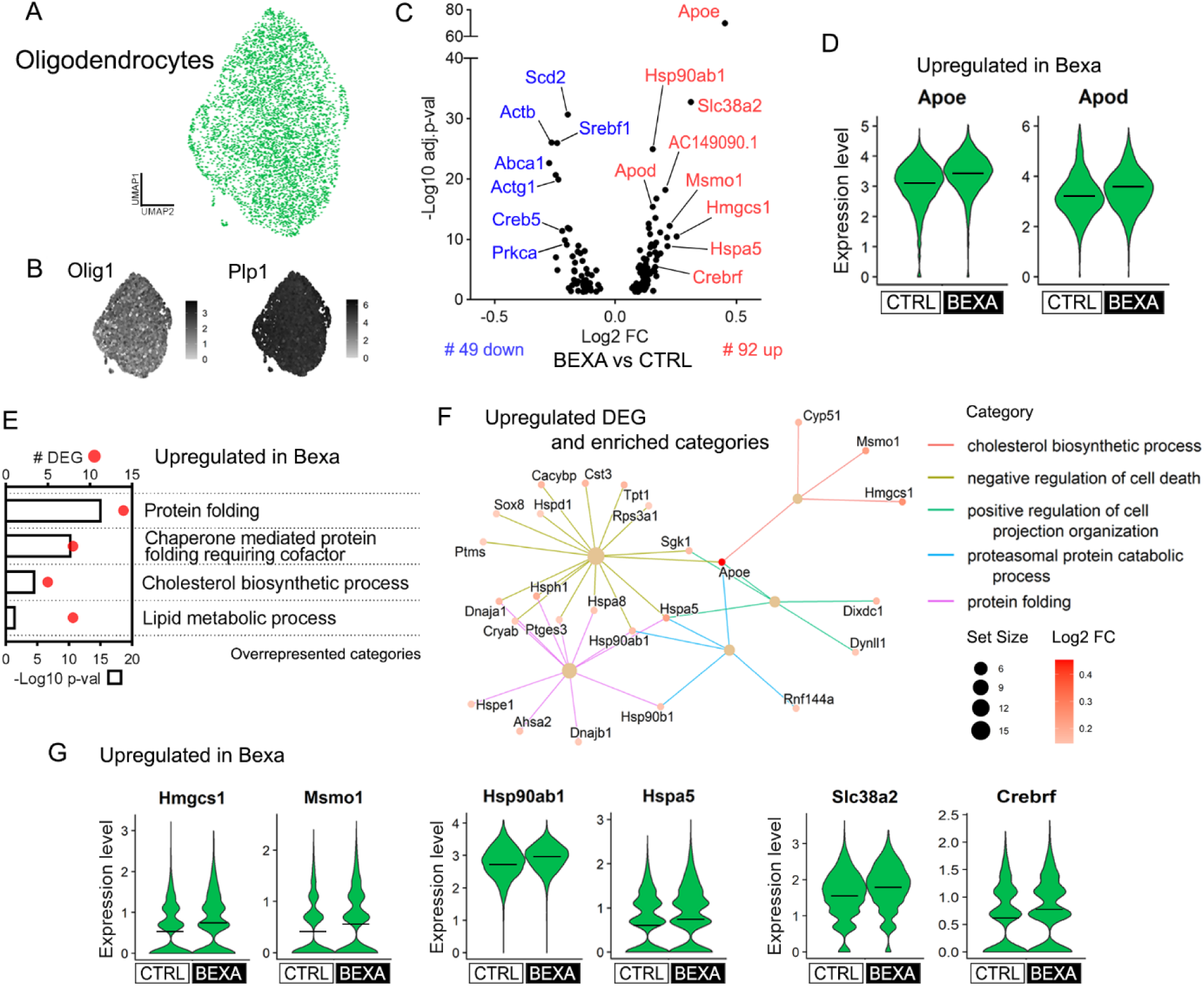
Oligodendrocyte gene expression responses to bexarotene treatment in APP/PS1 mice. (A) UMAP plot of 4,987 cells with oligodendrocyte gene expression profile. (B) Feature plots of gene markers: Olig1 (oligodendrocyte lineage) and Plp1 (mature oligodendrocytes). (C) Bexa vs Ctrl DE analysis found 141 DEG within oligodendrocytes, and the Volcano plot displays the 79 upregulated and 45 downregulated DEG. (D) Expression level of upregulated Apoe, Apod, in Bexa oligodendrocytes. (E) GO terms (biological process) associated with the list of log2FC>0.1 upregulated genes. Bars indicate -log10 p-val and dots indicate DEG count. (F) Category netplot of upregulated DEG in Bexa group, connected with the categories found in GSEA analysis. Scales indicate the gene set size and DEG log2FC. (F) Expression level of upregulated genes linked to Cholesterol biosynthetic process (Hmgcs1, Msmo1), Protein folding (Hsp90ab1, Hspa5), and CREB pathway (Slc38a2, Crebrf) in Bexa within oligodendrocytes. Crossbars indicate mean values. Expression levels are displayed as SCTransform corrected UMI counts. Sample size: Bexa = 2,511, Ctrl = 2,476.

The terms Cholesterol biosynthetic process and Lipid metabolic process were activated in Bexa oligodendrocytes, exemplified by the upregulated DEG of the mevalonate pathway (*Hmgcs1*, *Msmo1*, *Idi1*, and *Cyp51*) (Figure 4E-G). Previous studies show that cholesterol synthesis is also essential for myelin sheath formation (61, 62). Among the 45 downregulated genes, we identified other fundamental controllers of lipid metabolism: *Abca1, Scd2* (Stearoyl-CoA Desaturase 2), and *Srebf1* (Sterol Regulatory Element Binding Transcription Factor 1) (Figure 4C). These genes respectively control cholesterol efflux, fatty acid biosynthesis, and lipogenesis – all consensus LXR target genes. Their downregulation suggests a suppression of key LXR-mediated responses in oligodendrocytes exposed to bexarotene.

Animal models of AD exhibit increased endoplasmic reticulum (ER) UPR stress and oxidative stress (5), which can also be regulated by RXR activation (63). In oligodendrocytes, many bexarotene-upregulated genes were linked to the term Protein folding, including multiple Heat Shock Protein (HSP) family genes (*Hsp90ab1*, *Hspa5*, *Hsph1*, *Hspa1b*, *Hspa1a*) (Figure 4E-G). HSP genes were also included in the upregulated Negative regulation of cell death (Figure 4F). These responses are crucial for maintaining cellular proteostasis and promoting survival under Aβ-induced ER stress. Additionally, they are closely linked to cholesterol metabolism in oligodendrocytes (58). Bexarotene treatment also affected the expression of genes involved in the CREB signaling pathways, which regulates cell growth and survival in response to stress, growth factors, and metabolic states. Bexarotene activated *Crebrf* (CREB3 Regulatory Factor) as well as cAMP/PKA/CREB transcriptional targets such as *Slc38a2* and *Sgk1* (Figure 4G). In oligodendrocytes, PKA/CREB promotes myelination (64). In parallel, *Creb5* (CREB5) and *Prkca* (PKCα kinase) were downregulated, along with cell motility-associated genes (*Actb*, *Actg1*, Figure 4C).

Furthermore, the subset of COP/OPC cells did not show significant transcriptional changes following bexarotene exposure. However, when analyzing the entire oligodendrocyte lineage (including mature and immature cells), we identified DEG with similar trends in COP/OPC (Figure S4C). Some of these DEG are linked specifically to COP/OPC-associated changes in gene expression rather than mature oligodendrocytes, such as several developmental genes (*Hes5, Tafa1*, *Ncan*, *Ifi27, AC149090.1*). This reveals that bexarotene influences neurodevelopmental pathways within the oligodendrocyte lineage, potentially affecting immature cells.

Oligodendrocyte function is also tightly linked to neuronal health, and bexarotene has been shown to regulate neuronal genes and processes in AD mouse models (22, 23). Within neuronal clusters, we identified nine distinct subpopulations, including small groups of excitatory, inhibitory, and immature neuroblasts (Figure S4D-E) (30). However, the highly variable distribution of these subclusters across samples and the limited number of cells across subclusters (Figure S4F) prevented a robust evaluation of bexarotene’s effects on neuronal populations.

### Brain endothelial cells’ gene responses are linked to protein folding and vasculogenesis

AD also relates to vascular dysfunctions including brain hypoperfusion, barrier leakiness, and other blood-vessel cell vulnerabilities (6, 65). At last, single-cell transcriptomics enabled us to examine bexarotene-induced gene expression changes in mural (cluster 4) and endothelial cells (EC, clusters 0 and 11) *in vivo.* Analysis of mural cells comprising pericytes and vascular smooth muscle cells did not display DEG following bexarotene treatment (not shown). In parallel, EC subset yielding final 5,430 high-quality cells were re-clustered into two subpopulations: EC0-Capillary/Arteriole and EC1-Venule (Figure 5A). EC0 expressed both arteriole (*Arl15, Mgp, Gkn13*) and capillary (*Ivns1abp, Hmcn1*) markers, with upregulated terms linked to cell cycle and migration (Figure 5B and S5A). EC1 showed upregulated venule markers (*Vwf, Il1r1, Cfh*) and other genes linked to translation and response to hypoxia (Figure 5B and S5A). Since EC1 primarily originated from a single animal (Figure S5B), we focused our analysis on EC0-Capillary/Arteriole.

**Figure 5.**
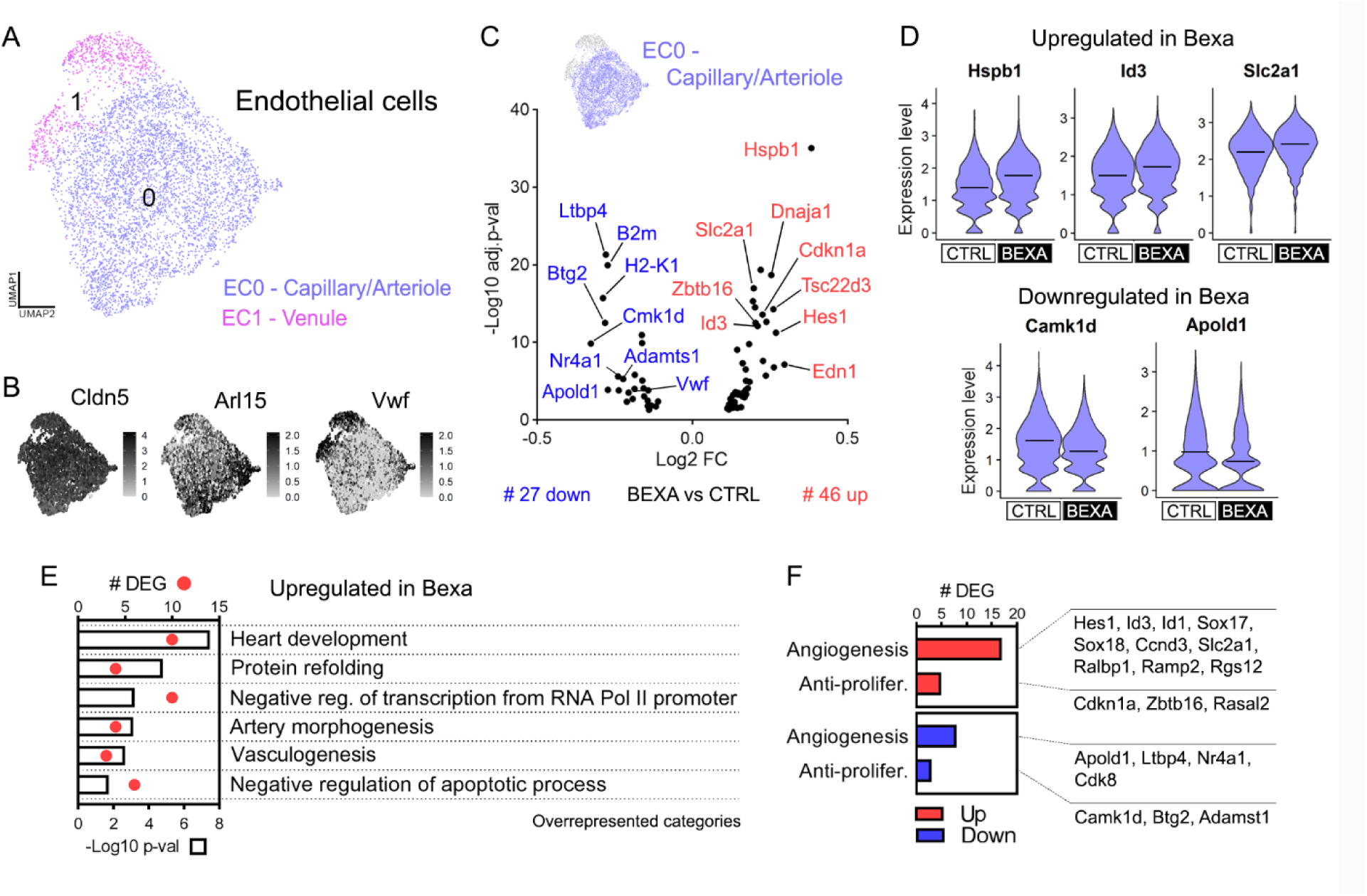
Brain EC molecular responses in APP/PS1 mice treated with bexarotene. (A) UMAP plot of the 5,430 cells with EC gene expression recovered from APP/PS1 mice brains, showing EC0 – Capillary/Arteriole and EC1 – Venule populations. (B) Feature plots of subclusters’ gene markers: Cldn5 (all EC), Arl15 (EC0, arteriole), and Vwf (EC1, venule). (C) Bexa vs Ctrl DE analysis found 73 DEG within the larger Capillary/Arteriole brain EC (EC0) population, and the Volcano plot displays the 46 upregulated and 27 downregulated DEG. (D) Expression level of upregulated Hspb1, Id3, and Slc2a1 in Bexa, and downregulated Camk1d and Apold1 within EC0 – Capillary/Arteriole. Crossbars indicate mean values, and the expression levels are displayed as SCTransform corrected UMI counts. (E) Overrepresented GO categories (biological process) associated with the list of log2FC>0.1 upregulated genes in Bexa group within EC0. Bars indicate -log10 p-val and dots indicate DEG count. (F) Counts for angiogenesis/EC proliferation marker genes upregulated (17) and downregulated (8) in Bexa group, and for anti-proliferative EC marker genes upregulated (5) and downregulated (3). Sample size: Bexa = 2,968, Ctrl = 1,707 (EC0).

Bexa vs. Ctrl DE analysis in EC0 identified 73 DEG, with *Hspb1* as the top upregulated gene (Figure 5C-D and Table S2). Similarly to oligodendrocytes, bexarotene increased the expression of HSPs and other protein refolding-associated genes (*Hspb1, Dnaja1, Hspa8, Hsp90ab1, Hsp90aa1*) (Figure 5E). Key upregulated GO terms included Heart development and Vasculogenesis, exemplified by the activation of *Hes1*, *Id3*, *Sox17* and *Sox18* (Figure 5D-E). The regulation of these and other transcription factors was reflected in the term Negative regulation of transcription from RNA Pol II promoter. Another top upregulated gene was *Slc2a1,* encoding the blood-brain barrier (BBB)-specific transporter GLUT1, which activates glucose uptake and EC proliferation (Figure 5D). GLUT1 levels are reduced in AD brain endothelium, contributing to neuronal loss and brain inflammation (6, 66). Other upregulated EC-associated genes included *Ctla2a, Pglyrp1, Edn1, Tbc1d4*, *Ramp2, Cavin2* and *Rgs12* (Table S1).

Bexarotene regulated genes involved in neuroinflammation control within ECs. RXR activation upregulated *Tsc22d3* (TSC22 Domain Family Member 3), a corticosteroid-responsive anti-inflammatory transcription factor, while downregulating *B2m* and *H2-K1* immune-associated genes in EC (Figure 5C and S5C). *Vwf* (Von Willebrand factor), which responds to vascular injury, tissue damage, and infection, was also downregulated in Bexa capillary/arteriole cells. VWF also modulates angiogenesis depending on its cellular localization (67).

Several Bexa-upregulated genes were linked to EC growth and apoptosis suppression (Figure 5E), though cell proliferation regulators showed both up- and down-regulated patterns (Figure 5F). In addition to GLUT1, HES, SOX, and Id genes, bexarotene-induced angiogenesis signals included *Ccnd3, Ralbp1*, *Rgs12*, and *Ramp2* (68–70). Conversely, anti-proliferative genes were downregulated, including *Camk1d* (CaM Kinase ID, Figure 5D)*, Btg2*, and *Adamts1.* However, cell cycle inhibitors were activated (*Cdkn1a, Zbtb16*) and EC proliferation marks such as *Apold1* (Apolipoprotein L Domain Containing 1, Figure 5D), *Ltbp4*, *Nr4a1,* and *Cdk8* were suppressed, thus indicating simultaneous signals for control of vascular cell growth. Protein refolding responses may also influence EC proliferation. HSPB1 mediates VEGF-induced EC cytoskeleton organization, with angiogenic effects varying by its intra- or extracellular localization (71). HSP90, encoded by the upregulated genes *Hsp90ab1* and *Hsp90aa1,* also controls EC growth by promoting cell proliferation and tube formation (72). As shown in Figure 5F, most pro-angiogenic markers were upregulated indicating an overall stimulus for brain capillary/arteriole growth following RXR activation.

### APOE-mediated signaling is a central axis of communication activated by bexarotene in brain cells

*Apoe*, a known RXR target gene, was upregulated in distinct cellular subsets following bexarotene treatment, as shown in Figure 6A. APOE interacts with multiple receptors to deliver lipids and lipoprotein-associated molecules, such as Aβ, to other cells. Activation of the APOE/TREM2 axis in reactive cells, for instance, is recognized as a key pathway for Aβ degradation and plaque compaction, triggering a downstream signaling cascade crucial for immune function (2). To predict the impact of bexarotene on cell signaling pathways, we next conducted network analysis of cell communication. Based on the gene expression profile of ligands, receptors, and cofactors in signaling pathways provided by ChellChatDB (37), we calculated and compared communication probabilities among cells acting as sources and targets of signals (Figure 6B and S6A). This analysis showed that bexarotene might strengthen the interactions between brain cells, particularly those involving astrocytes (Figure 6C and S6B-D). The number of significant interactions involving homeostatic microglia (Micro-Homeo) slightly decreased, likely reflecting the overall downregulation of gene expression induced by bexarotene in this population. The APOE-mediated signaling pathway appeared among the top signaling changes induced by bexarotene in the brains of APP/PS1 mice, in various cell types (Figure 6D). It showed an overall 30% increase compared to control, with 57% of the relative information flow linked to bexarotene network (Figure 6E and S6E). The main sources of APOE were Astro-Homeo, Astro-Reactive, Oligodendrocytes, and, to a lesser extent, ECs – all of which exhibited increased APOE signaling outgoing interaction strength. This pathway primarily targeted Micro-DAM and Micro-Homeo, both showing increased incoming interaction strength, as well as Astro-Reactive in the Bexa group (Figure 6E and S6E).

**Figure 6.**
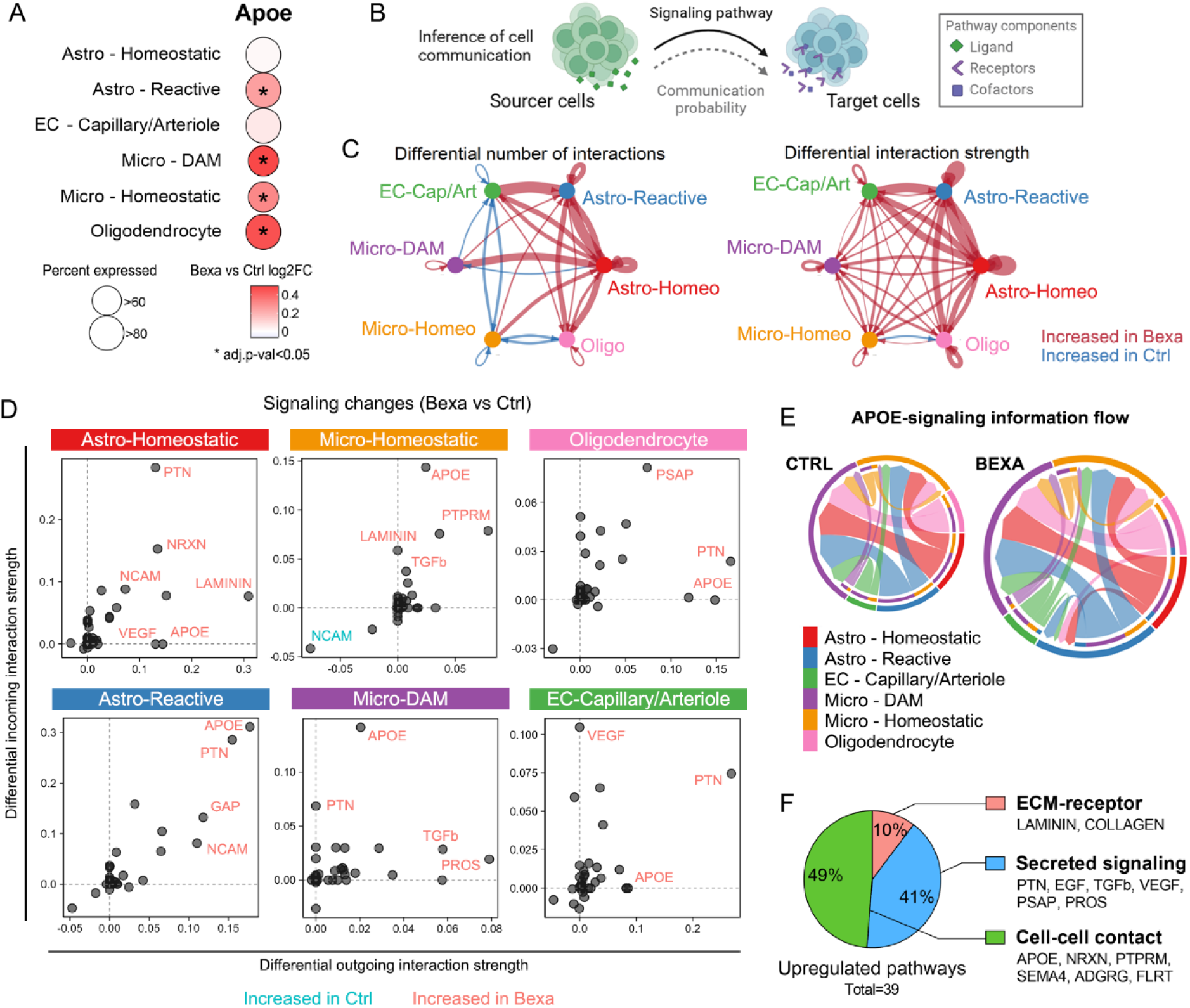
Cell communication inference reveals changes in APOE-mediated signaling by bexarotene. (A) Cell population-specific Apoe differential gene expression in response to the 10-day treatment with bexarotene in 5-month-old APP/PS1 mice. Circle size indicates the percent expressed in Bexa group. Color scale indicates Bexa vs Ctrl log2FC values. (B) CellChat was used to predict the impact of bexarotene-induced gene expression changes on the cell communication between populations. It was based on the expression profiles of gene products acting as ligands in the cell sources, and receptors/cofactors in the cell targets. (C) Circle plots showing the overall differential number of interactions (left panel) and differential interaction strength (right panel) among distinct cell populations, in Bexa compared to Ctrl group. Red arrows indicate increased interactions in Bexa, while blue arrows indicate increased interactions in Ctrl. (D) Signaling changes in Bexa vs Ctrl comparison for each cell population, showing that APOE appeared among the top axes of communication affected. The changes are expressed in terms of differential incoming (y axis) and outgoing (x axis) interaction strengths for each signaling pathway identified from CellChatDB. (E) Chord diagrams showing the APOE-signaling information flow in Ctrl and Bexa groups. The diagram size is proportional to the relative information flow, with 30% increase in Bexa relative to Ctrl. Outer bar colors represent the cell sources, and inner thinner bar colors represent the targets. Bar size is proportional to the signal strength sent or received. Edge colors are consistent with the sources, and edge weights are proportional to the interaction strength. (F) CellChatDB annotation of the signaling pathways strengthened by bexarotene. Among the 39 upregulated pathways (p-val<0.05), 10% are ECM-receptor interactions, 41% are secreted signaling pathways, and 49% are linked with cell-cell contact.

The Bexa group was associated with the upregulation of APOE and 38 other communication pathways (Figure S6E). These pathways were predominantly annotated as cell-cell contacts (49%) and secreted signalings (41%), with a smaller portion related to extracellular matrix (ECM)- receptor contacts (10%) (Figure 6F). Cell population-specific changes show that Astro-Homeo was central to many of those changes, supported by the strong DE among these cells (Figure 6D and S6F). For example, LAMININ pathway of ECM-receptor contact was increased between Astro-Homeo and other subsets, driven by the upregulation of *Lama3* (Laminin Subunit Alpha 3) in astrocytes (Figure S6F and Table S2). Neurexin-mediated contacts (NXRN) were upregulated between homeostatic astrocytes and other cells, associated with increased expression of pathway components like the receptor *Dag1* (Dystroglycan 1) in Astro-Homeo (Figure S6F and Table S2).

Signaling pathways coordinated by secreted growth factors were also increased linked to Astro-Homeo activity. PTN (Pleiotrophin) signaling, which promotes post-developmental neurotrophic and protective effects (73), had its interaction strength increased from but particularly towards homeostatic astrocytes, linked to increased levels of *Sdc4* (Syndecan 4) in these cells (Figure S6F and Table S2). Additionally, EGF (Endothelial Growth Factor) and VEGF signaling pathways were increased due to changes in the expression of their ligands in Astro-Homeo, connecting these cells with other astrocytes and EC, respectively (Figure S6F). These findings reflect the observed DE linked to neurodevelopment and angiogenesis. In contrast, analysis of signaling changes related to Astro-Reactive, showed minimal correspondence with significant changes in DE, likely due to the small number of cells in this subcluster.

In addition to APOE, pathways strengthened in the microglia populations included PTPRM (Protein Tyrosine Phosphatase Mu), SEMA4 (Semaphorin 4A), and TGFb (Transforming Growth Factor Beta), all linked to microglial cell growth and motility (Figure 6D). These changes were also connected to DE in homeostatic astrocytes, with the upregulation of *Ptprm*, semaphorin-coding genes, and *Tgfb2*. (Figure S6F and Table S2). For instance, the anti-inflammatory TGF-β cytokine is a key controller of microglia activation, and impairment of TGF-β pathway is a feature of AD-linked neuroinflammation (74). Notably, a previous study identified upregulated TGF-β levels in the brains of APP/PS1 mice treated with bexarotene (57). These predicted changes in cell communication, centered around APOE signaling, cell growth, and immune control, reflect the biological processes affected by bexarotene across brain cell subpopulations, and open new possibilities for studying RXR-regulated molecular targets and signaling pathways in AD physiopathology.

## Discussion

Recent single-cell approaches have revealed brain cellular heterogeneity in various contexts, including AD (49). However, how discrete cell populations respond to potential therapeutic interventions, such as NR-targeted treatments, remains largely unknown. We investigated bexarotene’s molecular impact across brain cells in APP/PS1 mice, generating a comprehensive set of DEG. Functional predictions, based on Gene Ontology (GO) terms and cell communication pathways, revealed that bexarotene-activated RXR receptors regulate cholesterol metabolism, development, cell growth, immune reactivity, and protein folding gene sets in a cell subpopulation-specific manner. Consistent with the present findings, transposase-accessible chromatin sequencing (ATAC-seq) profiles and integrative regulatory network analysis indicated that treatment led to differentially bound TFs linked to development in astrocytes and oligodendrocytes, angiogenesis in endothelial cells, and immune activation in microglia^1^.

*Apoe* activation was prominent in DAM and homeostatic microglia, oligodendrocytes, and reactive astrocytes of bexarotene-treated APP/PS1 mice. Our findings also show that *Apoe* upregulation is stronger in reactive and disease-associated glial cells compared to homeostatic cells. Increased APOE gene or protein expression has been previously described in the brains of APP/PS1 (18, 57) and APOE3 mice (75) following short-term RXR ligand treatment (up to 10 days). Similar effects were observed with long-term treatment in 24-month-old 3xTg-AD mice (13). LXR and PPARγ agonists also promote this effect in APP/PS1 mice, with simultaneous activation of both receptors exerting a synergistic effect on *Apoe* activation (21, 76). Here, we identify specific brain cell subpopulations that may mediate the *Apoe* upregulation linked to RXR activation. Moreover, we confirm the central role of APOE biology in the molecular effects of bexarotene in the brains of APP/PS1 mice, with the APOE-mediated signaling pathway being one of the top upregulated communication axes in the bexarotene group, primarily linking homeostatic and reactive glial cells. Bexarotene-mediated Aβ clearance and cognitive impairment seem to require functional APOE *in vivo* (9), and clinical evidence regarding bexarotene in AD shows increased serum Aβ and reduced CSF Aβ selectively in APOE4 non-carrier AD subjects treated with bexarotene (77). Therefore, the relationship between bexarotene signaling and APOE biology is critical for the potential clinical use of RXR ligands in AD.

The influence on APOE lipidation status through ABCA1 and ABCG1 is also considered a key aspect of the neuroprotective effects of bexarotene in AD models (15, 75). We and others have shown the upregulation of *Abca1/Abcg1* in the brains of bexarotene-treated AD mice, controlled by LXR/RXR (13, 18, 78). However, our results indicate that *Abca1/Abcg1* was either unaffected or downregulated by bexarotene in specific subpopulations, suggesting that this LXR-controlled response may vary depending on cell types and/or the physiological context. Previous studies using bulk RNA-seq of whole brains might have captured *Abca1/Abcg1* upregulation linked to cell populations not adequately represented by scRNA-seq, such as large and morphologically complex cells. Additionally, our findings reflect early-stage amyloid pathology, unlike previous studies that used older animals and different AD models. Notably, RXR-induced *Abca1* upregulation occurs only under high Aβ levels (78), indicating that RXR control over *Abca1/Abcg1* may vary according to model and disease stage. Given the importance of APOE lipidation in AD pathophysiology, further studies are needed to better understand the roles of ABCA1 and ABCG1 in RXR-mediated brain responses across cell subtypes and disease stages.

LXR/RXR, PPAR/RXR, and RAR/RXR play roles in the brain metabolic dysfunctions associated with AD (9, 75, 79). Thus, bexarotene’s metabolic effects reflect a complex network of NR signaling pathways, likely combining synergistic and conflicting responses. Our findings show that bexarotene significantly activates cholesterol and lipid biosynthesis pathways in homeostatic astrocytes, indicating an increase in astrocyte-derived cholesterol metabolism. Cholesterol synthesis and APOE-mediated transport are essential for brain repair (4). Cholesterol delivered to other brain cells influences neuronal neurotransmitter release, synaptic activity, oligodendrocyte myelination, and OPC proliferation and migration (3). The upregulated cholesterol synthesis pathway in oligodendrocytes, distinct from that in astrocytes, may be tailored to meet local demands (80). The regulation of cholesterol synthesis, CREB signaling, and *Apoe/Apod* expression in oligodendrocytes induced by bexarotene may support brain remyelination, especially at advanced stages of neuronal and myelin loss (58, 60, 81). Therefore, the activation of lipid metabolism in APP/PS1 astrocytes and oligodendrocytes may be connected to the observed neurorepair following brain RXR activation (9, 10).

As in our previous bulk RNA-seq study (18), single-cell transcriptional profiles of bexarotene-treated APP/PS1 mice revealed modulation of immune-associated gene responses in brain cells. Bexarotene activated immunoreactivity genes in reactive astrocytes and the classical DAM subpopulation, while downregulating DAM- and plaque-associated genes in homeostatic populations. In parallel, ECs showed downregulated immune activity. These findings highlight the selective enhancement of immune features in reactive cells. Bexarotene-induced *Apoe* upregulation was most prominent in DAM microglia, where APOE is a key feature of DAM states linked to the engulfment of apoptotic bodies and myelin debris (48). APOE also regulates microglial interaction with Aβ and plays a crucial role in plaque compaction (55, 56). Astrocytes, in an APOE-dependent manner, contribute to Aβ degradation, and deficits in astroglial Aβ clearance are implicated in AD pathogenesis (45, 46). The putative increase in Bexa-associated APOE signaling suggests activated Aβ processing and downstream cell activation through the APOE/TREM2 axis. Notably, bexarotene enhances immune cell phagocytic activity (18). Additionally, APOE-dependent regulation of cholesterol in lipid rafts links to its immunomodulatory roles. In activated microglia, cholesterol enrichment enhances phagocytosis and the release of neurotrophic factors (54). Thus, the combination of high *Apoe* and low *Abca1/Abcg1* observed especially in DAM phenotype may contribute to immune signaling transduction.

Previous studies have shown reduced neuroinflammation following RXR activation in AD models (12), consistent with our findings. Bexarotene may regulate immune reactivity also by upregulating anti-inflammatory cytokine genes like TGF-β, IL-6, and CCL2, while reducing IL-1β (57). Here, we further observed the increase in TGF-β pathway especially from homeostatic astrocytes towards microglia. PPAR activation antagonizes NF-κB signaling through TGF-β production (41), which may explain bexarotene’s effects. These findings highlight RXR activation’s dual role in immune responses: enhancing APOE-dependent DAM responses while reducing neuroinflammation.

These immune-related events were accompanied by decreased oxidative stress and injury responses in astrocytes and microglia, while protein folding and HSP-associated responses were increased in oligodendrocytes and ECs. HSP activation occurs in response to UPR to resolve ER stress, maintain proteostasis, and control cytotoxicity (82). Moreover, we and others have demonstrated that extracellular HSPs may promote neuroprotection and reduce neuroinflammation (83, 84). Notably, RXR activation has been demonstrated to regulate Aβ-induced ER stress in a neuroprotective manner (63). Protein folding responses are also tightly connected to PPARγ and LXR signaling and lipogenesis (85, 86), which may link to the effects observed in oligodendrocytes. In ECs, it is also connected to cell growth and vasculogenesis (71, 72).

Finally, we show that bexarotene activates developmental pathways especially in astroglia, but also in oligodendrocyte lineage cells and brain EC. Various RXR-containing dimers regulate cell growth, survival, and differentiation, and these pathways likely contribute to bexarotene-induced brain regeneration. Bulk RNA-seq revealed that bexarotene treatment activates brain development-associated genome events in AD mice brains (22, 23). Additionally, it promotes embryonic stem cells’ neuronal differentiation and enhances proliferation in neurogenic sites of APOE4 mice brain (23). *In vitro*, bexarotene promotes self-renewal proliferation and cell motility in adult NSCs, alongside improved late neuronal differentiation (24). Upregulation of SOX, HES, and Id family genes was observed in the single-cell profiles, along with subcluster-specific cell growth marks. For instance, the upregulation of genes like *AC149090.1*, within astrocytes and oligodendrocytes, may act linking metabolic responses to brain development (43). The Bexa-upregulated *Sox, Hes*, and *Id* regulate adult neurogenesis at multiple levels, usually by blocking the activation of proneural factors and maintaining self-renewal in progenitors (87). In glial differentiation, SOX2-deficiency in mature astrocytes compromises cell maturation and glutamate uptake (88). Thus, the activation of these features in astroglia could explain increased astrogenesis observed *in vitro* following bexarotene treatment (24). In oligodendrocytes, the upregulation of neurokinins and developmental genes, combined with changes in lipid metabolism and CREB pathway, aligns with prior observations of remyelination activation in bexarotene-treated AD mice (89). Additionally, RXRγ and 9-cis-RA are linked to OPC proliferation, differentiation, and myelination (25). Although we did not achieve sufficient resolution to identify OPC-specific responses to bexarotene, future studies could address this, especially in AD and other demyelination-related conditions.

The upregulation in *Sox, Hes*, and *Id* family genes was also observed in capillary/arteriole EC, with associated modulation of genes linked to vasculogenesis and increased VEGF signaling. In AD, ECs exhibit apoptosis and senescence, but these effects can be reversed by EC growth stimuli (90). Similarly, a human brain vascular atlas of AD revealed downregulated angiogenesis, proliferation, and cell motility in capillary and arterial EC (91). However, excessive VEGF signaling and non-productive angiogenesis, often observed in AD, contribute to the vascular imbalance (65). Therefore, we demonstrate that brain activation of RXR may support developmental pathways and cell growth also within EC, but the significance of such angiogenesis-related effect in AD needs further investigation.

## Conclusion

Our results provide a comprehensive overview of the molecular responses to RXR activation by bexarotene across various brain cell contexts. Bexarotene-treated APP/PS1 mice exhibit transcriptional activation of cholesterol biosynthesis and lipid metabolism in homeostatic astrocytes and oligodendrocytes; neurodevelopment in homeostatic astrocytes and oligodendrocyte lineage; cell growth and heat shock response in oligodendrocytes and endothelium; and *Apoe* activation and immune reactivity in reactive cells paralleled by a reduction in stress responses in homeostatic cells. Considering that therapies targeting multiple aspects of AD pathology may be required for optimal therapeutic efficacy, our findings thus highlight RXR and APOE biology as potential tools for developing therapies to modulate diverse biological processes in specific brain cells.

## Availability of data and materials

Raw data single-cell RNA-seq was submitted to GEO (https://www.ncbi.nlm.nih.gov/geo/, and will be publicly available as of the date of publication. This paper does not report the original code. Any additional information required to reanalyze the data reported in this paper is available from the corresponding author upon request.

## Abbreviations

9-cis-RA: 9-cis-retinoic acid
ABCA1: ATP-binding cassette transporter A1
ABCG1: ATP-binding cassette transporter G1
AD: Alzheimer’s disease
APOE: Apolipoprotein E
APP/PS1: APP/PS1ΔE9 transgenic mice
Aβ: Amyloid β
BBB: Blood-brain barrier
Bexa: Bexarotene
COP: Differentiation-committed oligodendrocyte progenitor
Ctrl: Control
DAM: Disease-associated microglia
DE: Differential expression
DEG: Differentially expressed gene
EC: Endothelial cell
ECM: Extracellular matrix
ER: Endoplasmic reticulum
FA: Fatty acid
HSP: Heat Shock Protein
LXR: Liver X Receptor
NR: Nuclear Receptor
NSC: Neural Stem Cell
OPC: Oligodendrocyte precursor cell
PC: Principal Component
PPAR: Peroxisome Proliferator-Activated Receptors
RAR: Retinoic Acid Receptor
RXR: Retinoid X Receptor
scRNA-seq: Single-cell RNA Sequencing
TF: Transcription Factor
UMAP: Uniform Manifold Approximation and Projection
UPR: Unfolded Protein Response

## Acknowledgments

The authors thank Mikayla Ranae McGuire and Mary Ann Ostach for their technical support.

## Authors’ contributions

Conceptualization, R.K., I.L. N.F.F, C.S-G; Methodology, C.S., Y.L., N.F.F.; Investigation, C.S-G, Y.L., N.F.F.; Formal analysis, C.S-G.; Visualization, C.S-G.; Writing – Original draft, C.S-G.; Writing – Review & Editing, C.S-G., R.K., I.L., D.P.G.; Supervision, R.K., I.L., D.P.G.; Project administration, R.K., I.L.; Resources, R.K., I.L.; Funding acquisition R.K., I.L.

## Ethics declarations

### Ethics approval and consent to participate

All animal procedures were performed according to the Guide for the Care and Use of Laboratory Animals from the United States Department of Health and Human Services, and were approved by the University of Pittsburgh Institutional Animal Care and Use Committee.

### Consent for publication

Not applicable.

## Supplementary information

### Supplementary figures

Figure S1. scRNA-seq data of bexarotene-treated APP/PS1 mice brains (Supplementary to Figure 1).

Figure S2. Astrocyte gene expression in Ctrl and Bexa (Supplementary to Figure 2).

Figure S3. Microglia gene expression in Ctrl and Bexa (Supplementary to Figure 3).

Figure S4. Oligodendrocyte cell lineage and neuron gene expression in Ctrl and Bexa (Supplementary to Figure 4).

Figure S5. Brain EC gene expression in Ctrl and Bexa groups (Supplementary to Figure 5).

Figure S6. Prediction of bexarotene impact on cell communication networks (Supplementary to Figure 6).

Supplemental table 1

Information on mice treatment, single-cell sample preparation, and cell type gene sets used in the analysis.

Supplemental table 2

Cell subtype-specific DE analysis (Bexa vs Ctrl), containing the DEG list, average log2FC, fraction of Bexa-expressing cells (pct.1), fraction of Ctrl-expressing cells (pct.2), and adj. p-value.

## Footnotes

1 Manuscript under review.

