## Supplemental Figures for "Brain single-cell transcriptional responses to bexarotene-activated RXR in Alzheimer’s disease model"

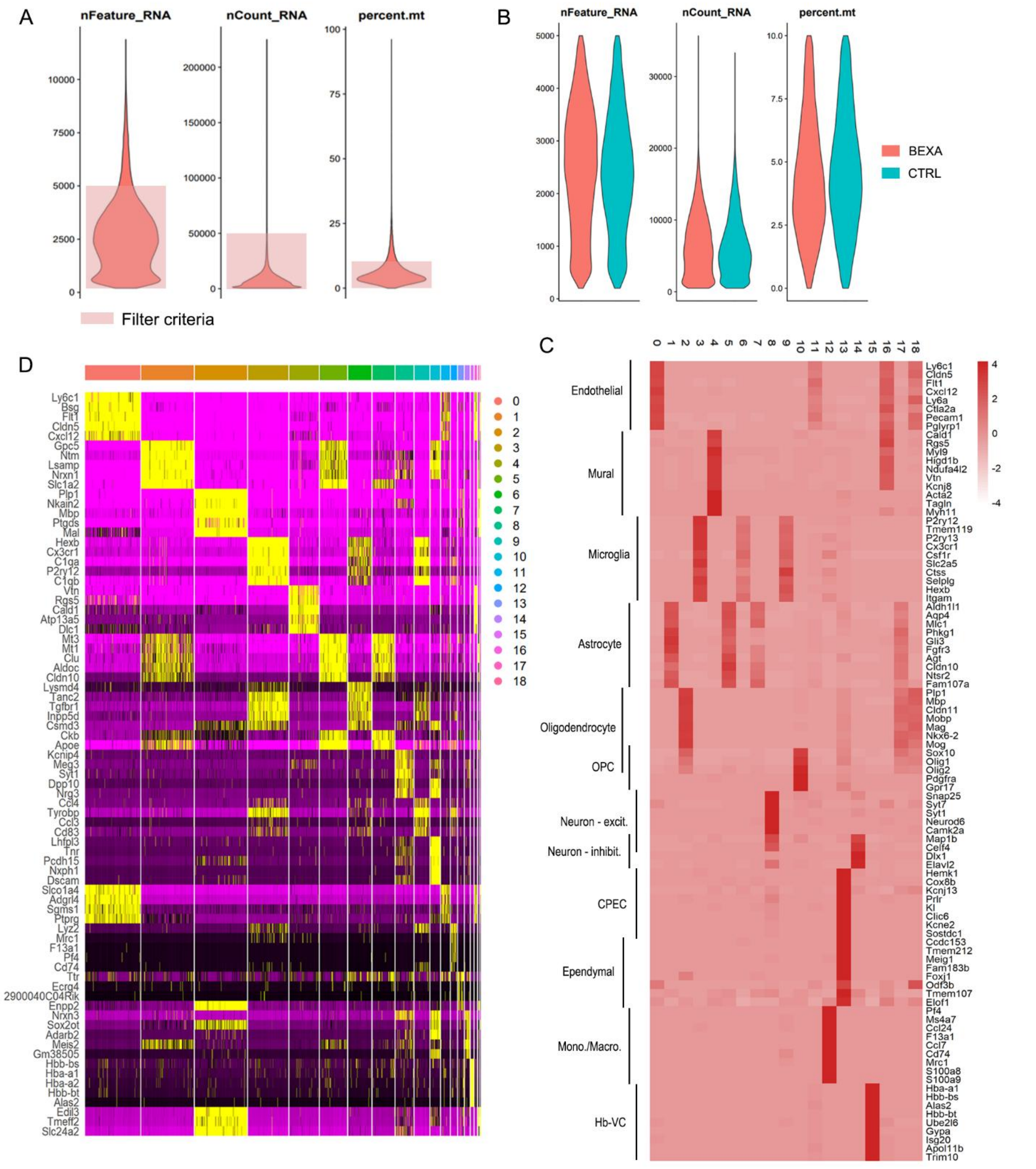


Supplemental Figure 1. scRNA-seq data of bexarotene-treated APP/PS1 mice brains (Supplementary to Figure 1). (A) Parameters of barcode quality control relative to the raw dataset: unique feature counts per barcode (nFeature_RNA), total counts per barcode (nCount_RNA), percentage of mitochondrial gene counts per barcode (percent.mt). The raw dataset was filtered to keep the barcodes with nFeature_RNA > 200 and < 5000, nCount_RNA < 50000, and percent.mt < 10, resulting in a matrix with total 23,745 genes across 37,634 cells. (B) Parameters of barcode quality control relative to the filtered dataset split by treatment group. (C) Clustering analysis identified 19 discrete clusters (0-18), and differential expression analysis comparing the clusters revealed the markers for each population. The heatmap depicts the SCT-transformed UMI counts of clusters' marker genes for a subset of barcodes (cells) from each cluster. Yellow indicates the higher expression, and magenta indicates the lowest. (D) The 19 clusters were manually annotated based on the expression levels of cell-type specific gene sets. For each cluster, the heatmap depicts the average SCT-transformed UMI count (scale indicated) of each cell-type marker used for annotation. The cell-type identity is indicated.

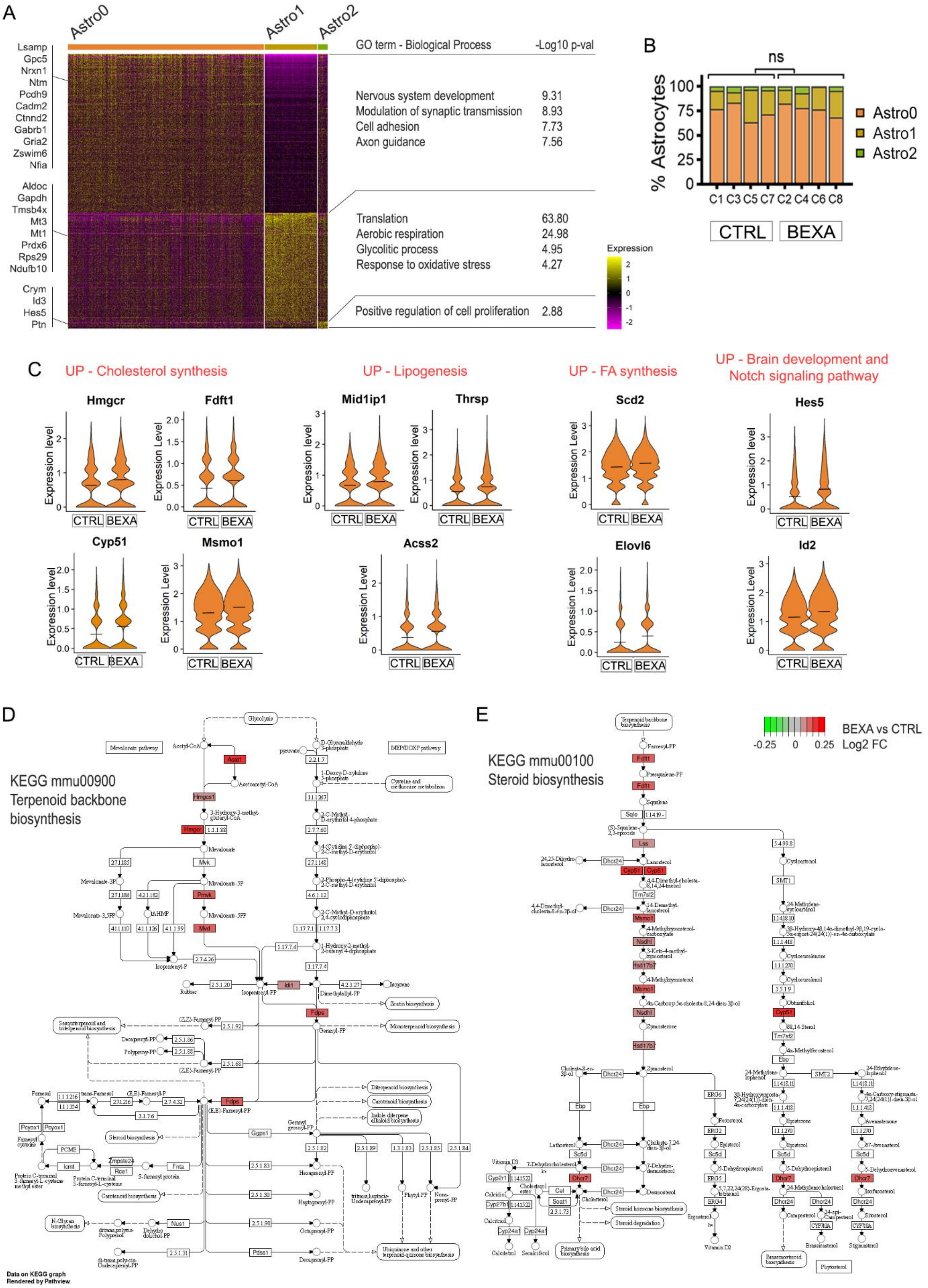
A
Astro0
Astro1
Astro2
Lsamp
Gpc5
Nrxn1
Ntm
Pcdh9
Cadm2
Ctnnd2
Gabrb1
Gria2
Zswim6
Nfia
Aldoc
Gapdh
Tmsb4x
Mt3
Mt1
Prdx6
Rps29
Ndufb10
Crym
Id3
Hes5
Ptn
GO term - Biological Process
-Log10 p-val
Nervous system development 9.31
Modulation of synaptic transmission 8.93
Cell adhesion 7.73
Axon guidance 7.56
Translation 63.80
Aerobic respiration 24.98
Glycolitic process 4.95
Response to oxidative stress 4.27
Positive regulation of cell proliferation 2.88
Expression
B
ns
% Astrocytes
Astro0
Astro1
Astro2
C1 C3 C5 C7 C2 C4 C6 C8
CTRL
BEXA
C
UP - Cholesterol synthesis
UP - Lipogenesis
UP - FA synthesis
UP - Brain development and Notch signaling pathway
Hmgcr
Fdft1
Mid1ip1
Thrsp
Scd2
Hes5
Cyp51
Msmo1
Acss2
Elovl6
Id2
Expression level
CTRL BEXA
D
KEGG mmu00900 Terpenoid backbone biosynthesis
Data on KEGG graph
Rendered by Pathview
E
KEGG mmu00100 Steroid biosynthesis
BEXA vs CTRL
-0.25 0 0.25 Log2 FC

Supplemental Figure 2. Astrocyte gene expression in Ctrl and Bexa (Supplementary to Figure 2). (A) Heatmap with top marker genes (log2FC>0.5) obtained for astrocyte subclusters after DE analysis. In the right, the GO terms linked to the markers lists after enrichment analysis (p-values indicated). (B) Distribution (%) of astrocyte subclusters within Ctrl and Bexa libraries. Two-way ANOVA statistical testing showed non-significant (ns) difference between groups: p-val>0.99, F=8.1×10-14. (C) Expression level (SCTransform corrected UMI counts) of Bexa-upregulated DEG within Astro0. Crossbars indicate mean values. (D-E) Pathview maps of Terpenoid backbone biosynthesis and Steroid biosynthesis KEGG pathways, respectively, with Bexa vs Ctrl DEG highlighted with log2FC values. DAVID enrichment analysis found -log10 p-values of 4.7 and 3.3 for each pathway, respectively.

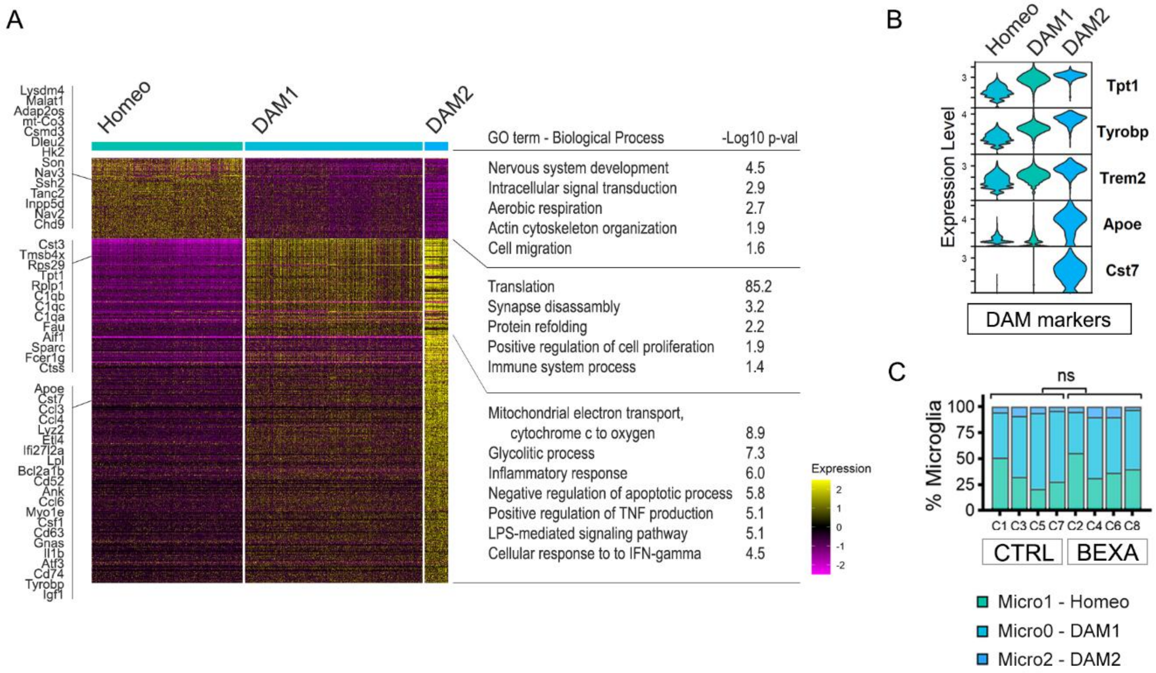


Supplemental Figure 3. Microglia gene expression in Ctrl and Bexa (Supplementary to Figure 3). (A) Heatmap with top marker genes (log2FC>0.5) obtained for microglia subclusters after DE analysis. In the right, the GO terms linked to the markers lists after enrichment analysis (p-values indicated). (B) Expression level of well-known DAM marker genes show DAM enrichment in Micro0 – DAM1 and Micro2 – DAM2. (C) Distribution (%) of microglia subclusters within Ctrl and Bexa libraries. Two-way ANOVA statistical testing showed non-significant (ns) difference between groups: p-val>0.99, F=2.4×10-32.

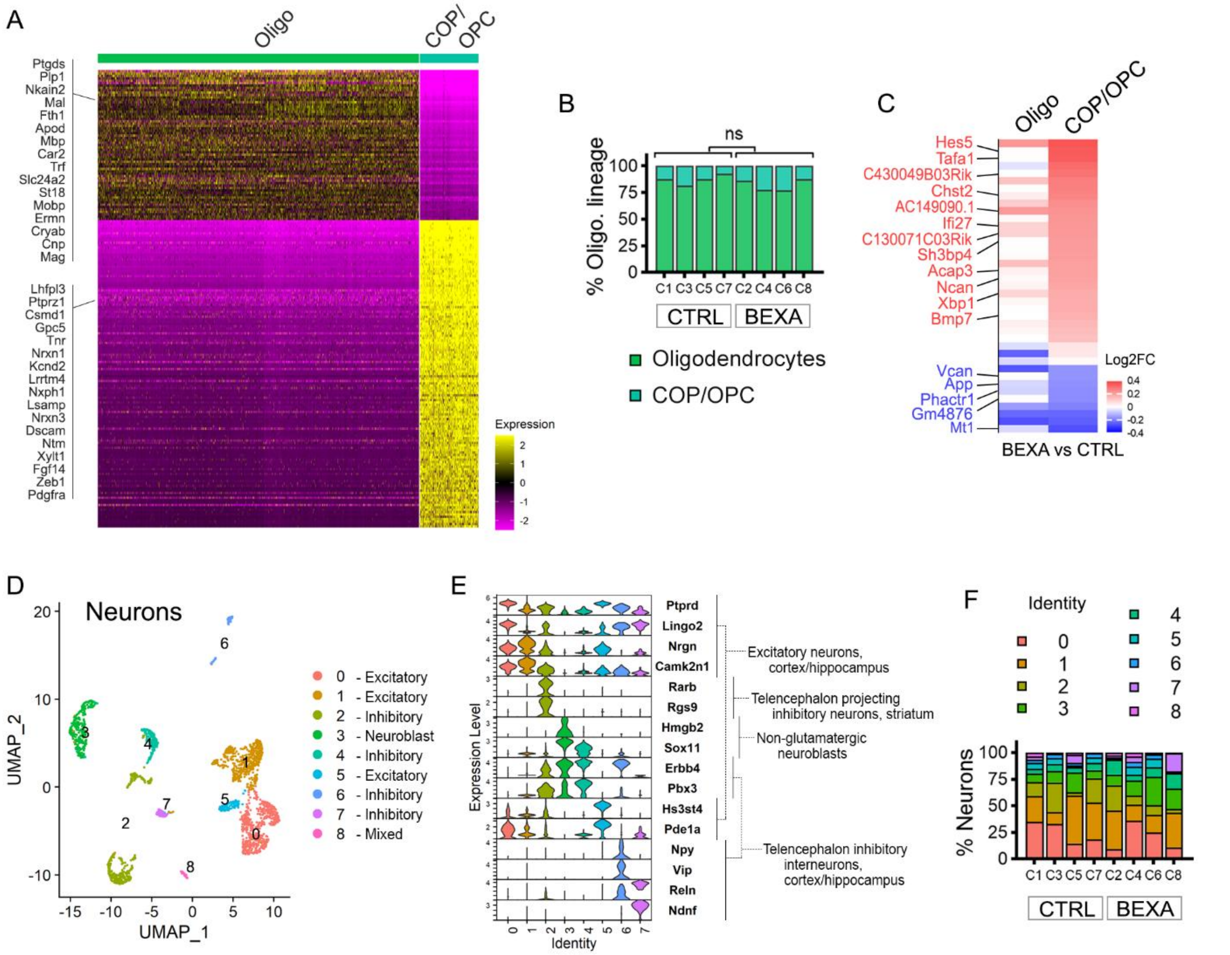


Supplemental Figure 4. Oligodendrocyte cell lineage and neuron gene expression in Ctrl and Bexa (Supplementary to Figure 4). (A) Heatmap with top marker genes (log2FC>2.0) obtained for DE analysis of clusters with oligodendrocyte lineage gene expression: Oligodendrocytes and COP/OPC. (B) Distribution (%) of oligodendrocyte cell lineage clusters within Ctrl and Bexa libraries. Two-way ANOVA statistical testing showed non-significant (ns) difference between groups: p-val>0.99, F=6.2×10-18. (C) Bexa vs Ctrl DE analysis considering all oligodendrocyte lineage population (Oligodendrocytes & COP/OPC) revealed a list of all Oligo DEG. The heatmap depicts the Bexa vs Ctrl log2FC of such DEG when each population is analyzed separately. (D) UMAP plot of neuronal subpopulations containing a total of 2,223 cells. (E) Expression level of subclusters' gene markers, with the respective neuronal identity indicated. Two markers are displayed for each subcluster and are ordered by subcluster number. (F) Distribution (%) of neuronal subclusters within Ctrl and Bexa libraries.

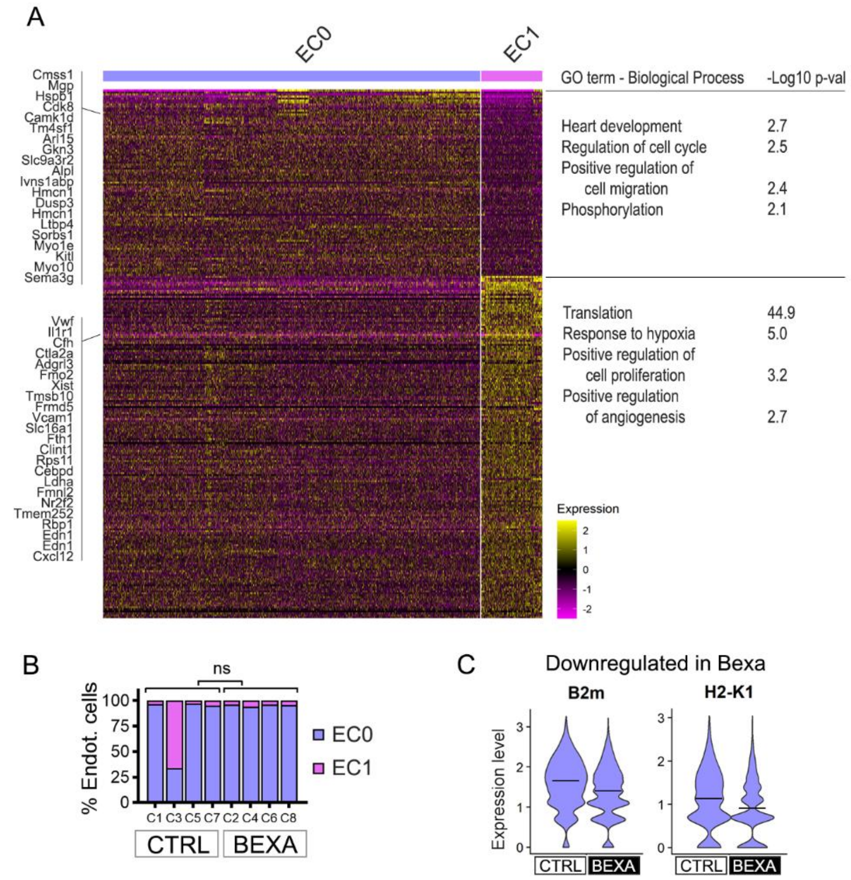


Supplemental Figure 5. Brain EC gene expression in Ctrl and Bexa groups (Supplementary to Figure 5). (A) Heatmap with top marker genes (log2FC>0.25) obtained for Endot subclusters after DE analysis. In the right, the overrepresented GO terms linked to the markers lists (p-values indicated). (B) Distribution (%) of EC subclusters within Ctrl and Bexa libraries. Two-way ANOVA statistical testing showed non-significant (ns) difference between groups: p-val=0.2909, F=1.221. (C) Expression level (SCTransform corrected UMI counts) within EC0 for Bexa and Ctrl group of downregulated genes linked to inflammation (*B2m, H2-K1*). Crossbars indicate mean values.

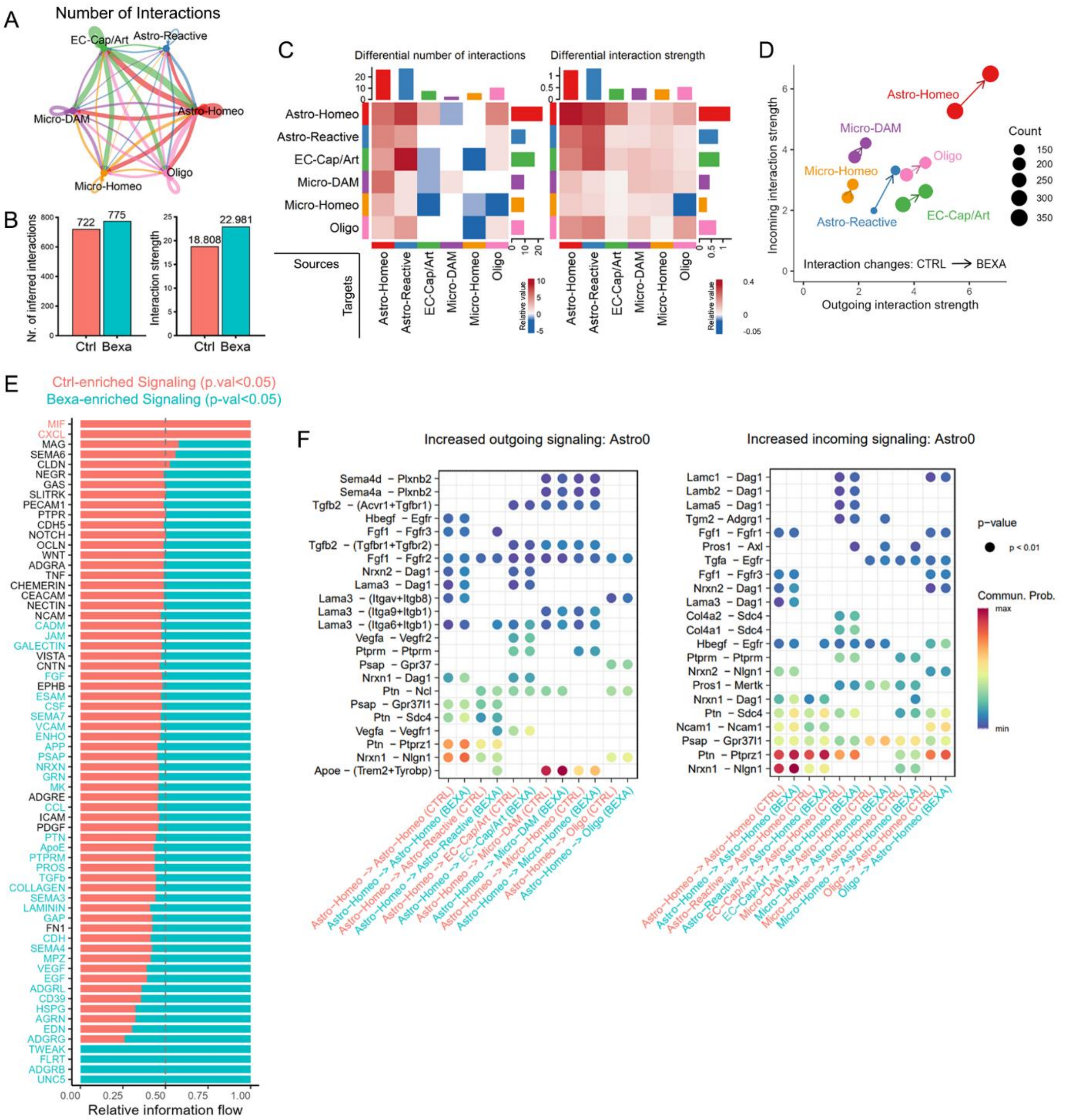


Supplemental Figure 6. Prediction of bexarotene impact on cell communication networks (Supplementary to Figure 6). (A) Circle plot of the aggregated network of cell communication computed based on gene expression profiles, using CellChat. Arrow thickness indicates the number of interactions from distinct cell populations in the dataset. (B) Comparison of the number of interactions and interaction strength, respectively, in Ctrl and Bexa groups. (C) Bexarotene-mediated changes in the number of interactions and interaction strength, respectively, among different cell populations. The sources (in each row) are the populations acting as origin of signaling pathways, and the targets (in each column) are the receiver of communication input. The top bar plot represents the sum of each column of the absolute values displayed in the heatmap (incoming signaling). The right colored bar plot represents the sum of each row of the absolute values (outgoing signaling). (D) Overall comparison of the incoming (y axis) and outgoing (x) interaction strengths for each cell population analyzed, in Bexa compared to Ctrl group. The circle size indicates the number of interactions in each group, and the arrows indicate the shift in the interaction strengths induced by Bexa. (E) Relative information flow between Ctrl (red bars) and Bexa (green bars) for each signaling pathway, considering all cells. Information flow: the sum of communication probability among all pairs of cell groups. Signaling pathways were ranked based on the differences in the overall information flow. Ctrl- and Bexa-enriched signaling pathways are indicated, defined as the ones with significant differences in the flow of information between the two conditions (paired Wilcoxon test, p-val<0.05). (F) Bubble plots showing the communication probabilities of ligand-receptor (LR) pairs of pathways increased in Astro – Homeo (Astro0) due to bexarotene. CTRL and BEXA selective pairs of cell groups are displayed (x axis) for each LR pair (y axis). The left panel shows communication with Astro – Homeo as source (outgoing), and the right panel shows Astro – Homeo as target (incoming). Only significant LR-pairs (p-val<0.05) are shown.
